# A Grape Seed Oligomeric Procyanidin Extract Reverses Diet-Induced Obesity Through Gut Microbiota Remodeling and Restoration of GLP-1, Gut-Brain, and Gut-Liver Signaling

**DOI:** 10.64898/2026.07.31.741918

**Authors:** Mohamed Mokrani, Romain Villéger, Amira Guellim, Ludivine Nardy, Manoli Leclaircie, Lucie Lebeau, Marc Biran, Hélène Roumes, Amandine Brochot, Anne-Karine Bouzier-Sore, Maria C. Urdaci

## Abstract

**Background:** Obesity is a complex multifactorial disease associated with chronic low grade inflammation, gut microbiota dysbiosis, and impaired gut-brain communication. Oligomeric procyanidins from grape seed extracts (GSE) are promising prebiotic candidates, capable of modulating host metabolism through interactions with the gut microbiota.

**Methods:** C57Bl/6J male mice were rendered obese by feeding them a high fat, high sucrose diet and were orally administered GSE at a dose of 1or 2 g/kg/day for 12 weeks. We assessed body weight, adiposity, glucose tolerance, insulin sensitivity, circulating hormones, brain homeostasis markers, colonic and liver gene expression, 16S rRNA gene sequencing of the gut microbiota profiles, and untargeted cecal metabolomics.

**Results:** GSE reduced body weight gain, visceral adiposity, adipocyte hypertrophy, and improved oral glucose tolerance and insulin sensitivity. It normalized circulating lipid and glucose levels and lowered fasting insulin and leptin while increasing endogenous GLP-1. Hepatic gene expression analysis revealed a dose-dependent restoration of antioxidant defenses (SOD, CAT) and lipogenic transcription factors (SREBP, ChREBP). In the colon, GSE attenuated pro-inflammatory IL6 cytokine expression and strikingly upregulated GLP-1 and GLP-1 receptor expression. Microbiota analysis revealed a profound, dose-dependent remodeling of gut microbiota composition and diversity, with an expansion of health-associated taxa, such as *Akkermansia muciniphila*. Brain analyses revealed restoration of NAA and BDNF levels together with markers consistent with improved mitochondrial function. Cecal metabolomics revealed normalization of secondary bile acid metabolism, restoration of arginine bioavailability, and reduction in the accumulation of L-DOPA and spermidine.

**Conclusions:** An oligomeric procyanidin-rich grape seed extract acts as a multitarget prebiotic that alleviates diet-induced obesity and is associated with coordinated restoration of gut microbiota composition, GLP-1 signaling, and gut–brain and gut-liver communication pathways. Convergent dose-dependent effects on *Akkermansia muciniphila* abundance, GLP-1, NAA, and BDNF identify key mechanisms underlying its metabolic benefits.

## Introduction

Obesity represents one of the most consequential metabolic disorders of our time, affecting more than one billion adults worldwide and projected to impact nearly 60% of the global adult population by 2050 ^1^.

Beyond its epidemiological magnitude, obesity represents a profound disruption of systemic metabolic homeostasis, driving insulin resistance, ectopic lipid accumulation, non-alcoholic fatty liver disease (NAFLD), and chronic low-grade inflammation that underlies many of its major comorbidities, including type 2 diabetes, cardiovascular disease, and mental health ^2^. Alarmingly, obesity now affects a greater proportion of children and adolescents (5–19 years) worldwide than underweight, with unhealthy food environments identified as a major driver of the rapid increase in pediatric overweight and obesity ^3^. This escalating prevalence is expected to place an unprecedented burden on healthcare systems worldwide. By 2050, an estimated 3.8 billion adults worldwide, around 60% of the global adult population, are projected to be living with overweight or obesity ^1^. Therefore, a comprehensive understanding of the multifactorial mechanisms underlying obesity, together with the development of effective interventions spanning early-life prevention to strategies in adulthood, is essential to mitigate the growing global health and socioeconomic burden associated with this disease ^4^.

Obesity is a complex, multifactorial disease arising from the interplay between genetic predisposition, epigenetic modifications, environmental pressures, and behavioral factors that collectively disrupt energy homeostasis ^2^. Low-grade systemic chronic inflammation (SCI) is a crucial factor in the development of obesity and metabolic disorders. ^5,6^. The accumulation of pro-inflammatory chemokines and cytokines in tissues is considered a hallmark of SCI. It contributes to cardiovascular disease, cancer, diabetes mellitus, chronic kidney disease, NAFLD, and autoimmune and neurodegenerative disorders. ^5^. The gut microbiota has emerged as a central, causally implicated orchestrator of obesity. In diet-induced obesity, profound compositional and functional dysbiosis of the intestinal microbial community drives increased intestinal permeability, thereby promoting the translocation of microbial metabolites and activating innate immune pathways that sustain inflammation in hepatic and adipose tissues.^7,8^. Mechanistic studies of microbiota–host interactions have revealed remarkable specificity. For example, the microbiota-derived metabolite indole-3-propionic acid enhances mitochondrial respiration in CD4⁺ T cells, thereby suppressing intestinal inflammation^9^. Whereas microbially produced trimethylamine attenuates metabolic inflammation through the targeted inhibition of IRAK4, a key kinase in the innate immune cascade ^10^. Collectively, these findings establish that gut microbial metabolites are not merely associated with metabolic dysfunction but actively regulate i host immune and metabolic programs through defined molecular mechanisms.

Two major bidirectional communication axes critically connect intestinal microbial function with systemic homeostasis. Along the gut-liver axis, dysbiosis-driven impairment of intestinal barrier integrity compromises the physiological trafficking of microbial metabolites, bile acid signaling, and hepatic lipid and glucose metabolism, thereby promoting the development of NAFLD and hepatic insulin resistance ^11,12^. In parallel, the gut microbiota exerts profound regulatory effects on central nervous system function through the gut-brain axis ^13^.

Along the gut-brain axis, the gut microbiota regulates enteroendocrine secretion, neurotransmitter biosynthesis, glial cell function, and neuropeptide signaling, including the release of glucagon-like peptide-1 (GLP-1), a critical incretin with potent insulinotropic, anorexigenic, and neuroprotective properties ^13,14^. The pharmacological exploitation of the GLP-1 signaling axis through receptor agonists has established transformative efficacy in obesity and type 2 diabetes management, including sustained cardiovascular protection in large-scale randomized trials ^15–20^. Nevertheless, GLP-1 receptor agonists do not correct the underlying dysbiotic intestinal environment that contributes to metabolic disease. Consequently, identifying microbiota-targeting strategies that enhance endogenous GLP-1 secretion represents a promising, largely unexplored avenue. Beyond GLP-1, N-acetylaspartate (NAA), a highly abundant neuronal metabolite, is involved in metabolism and axonal myelination. NAA levels are significantly reduced in obesity and type 2 diabetes and are increasingly recognized as a marker of impaired neurometabolic homeostasis ^21^. Collectively, these findings support the concept that metabolic alterations in the gut and brain are tightly interconnected, reinforcing the concept of a bidirectional gut–brain regulatory network.

Polyphenols are plant-derived secondary metabolites that play a critical role in plant defense and survival, conferring protection against environmental stressors, including UV radiation and pathogenic microorganisms, through their antimicrobial and antioxidant activities. ^22–24^. Oligomeric procyanidins (OPCs) constitute a class of condensed tannins built from flavan-3-ol units, primarily catechin and epicatechin, interconnected through A-type or B-type interflavan linkages. ^25,26^. In a previous study, we demonstrated the anti-inflammatory, antioxidant, and antibacterial effects of grape seed extract (GSE) against selected pathogenic microorganisms ^24,25^. OPCs can be considered prebiotic compounds that positively modulate gut microbial communities. ^27–29^. Beyond their well-established antioxidant and anti-inflammatory properties within the gastrointestinal tract,,OPCs may exert broader systemic effects by modulating the gut-liver and gut-brain axis, both of which play central roles in metabolic regulation and neurological function. ^30–34^.

In a previous study, we demonstrated that GSE modulates key gut-derived signaling peptides, including GLP-1 and NPY, highlighting the potential of targeting gut-brain communication through dietary bioactive compounds. ^25^. Building on these findings, the present study aimed to evaluate the effect of the same OPC-rich GSE in a mouse model of diet-induced obesity, with particular emphasis on its prebiotic properties and its impact on gut microbiota composition, gut–brain signaling, and gut–liver communication pathways.

## Results

### GSE dose-dependently reduces body weight, tissue mass, and systemic metabolic disturbances

Twelve weeks of HFHS feeding produced robust obesity in C57BL/6J male mice, evidenced by a significant increase in cumulative body weight gain relative to standard-chow control group (CRL; ****p < 0.0001; Fig. 1A). The oral gavage with OPCs-rich GSE at both doses attenuated this body weigh gain: GSE1X (1 g/kg/day) reduced significantly body weight gain versus HFHS (***p < 0.001), and GSE2X (2 g/kg/day) prevent body weight gain (****p < 0.0001). At the same time, GSE1X animals remained modestly but significantly heavier than CRL mice (*p < 0.05). Tissue-level analyses confirmed dose-dependent adipose remodeling (Fig. 1B). HFHS markedly expanded eWAT mass (****p < 0.0001 vs CRL), and both GSE1X (*p < 0.05) and GSE2X (***p < 0.001) significantly reduced eWAT weight, with GSE2X achieving the greater reduction. BAT mass was paradoxically elevated by HFHS (****p < 0.0001 vs CRL), consistent with hypertrophic thermogenic dysfunction. GSE2X decreased BAT compared to the HFHS group (**p < 0.01 vs HFHS) and even normalized BAT compared to CRL. Hepatomegaly was induced in HFHS mice (*p < 0.05 vs CRL), and both GSE doses fully decreased liver weight to CRL level (**p < 0.01 each). Histological quantification of epididymal adipocyte numbers corroborated these findings: HFHS induced profound adipocyte hyperplasia (****p < 0.0001 vs CRL), whereas GSE1X and GSE2X (**p < 0.01) both significantly reduced adipocyte number (Fig. 1D).

**Figure 1.**
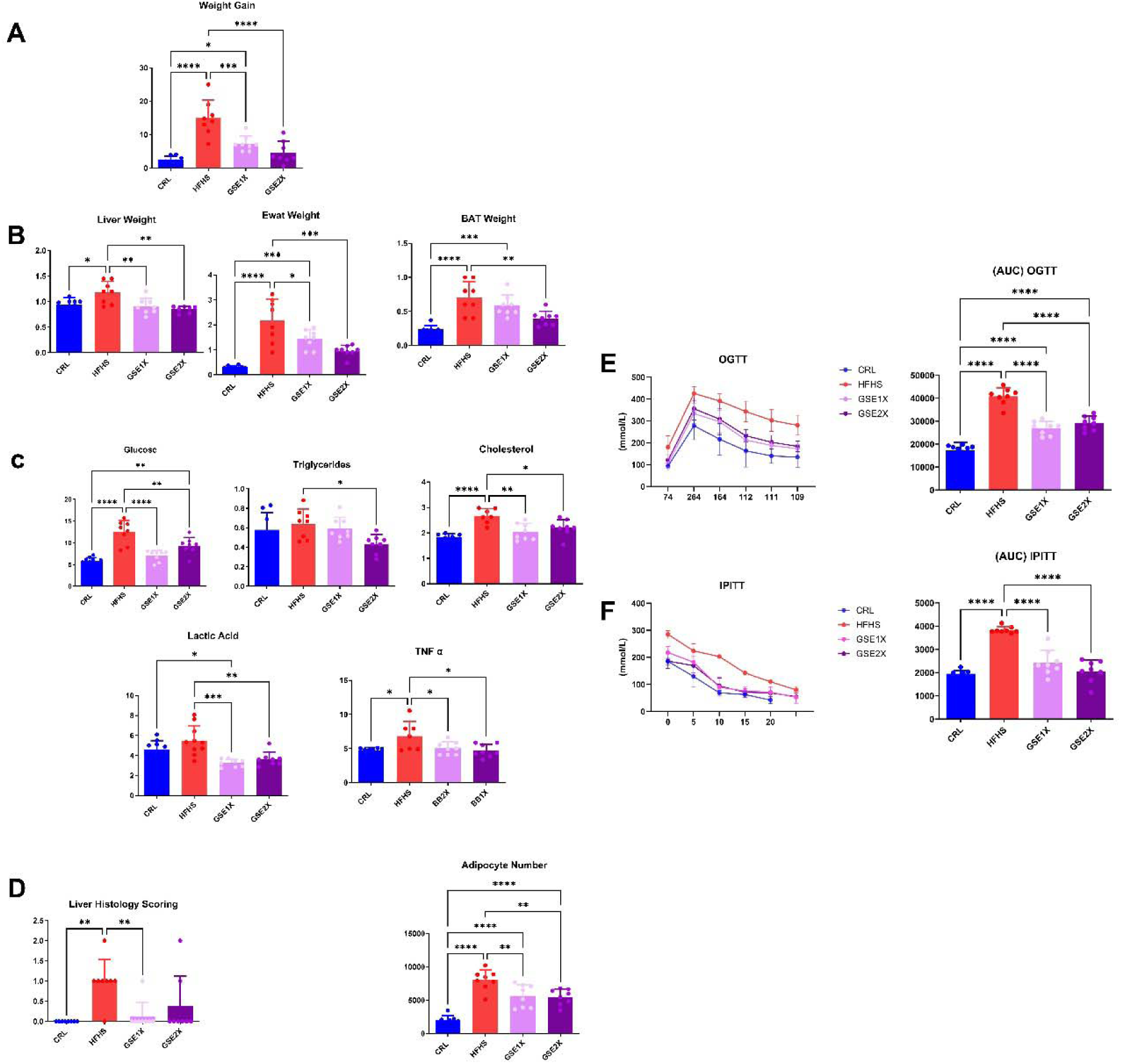
| GSE administation attenuates diet-induced obesity, metabolic tissue remodeling, fasting blood parameters, adipocyte hyperplasia and improves glucose tolerance and insulin sensitivity. (A) Total body weight gain (g) over the 10-week dietary intervention in CRL (standard chow), HFHS (high-fat high-sugar diet), GSE1X (HFHS + 1 g/kg/day GSE), and GSE2X (HFHS + 2 g/kg/day GSE) groups. (B) Post-mortem weights (g) of liver, epididymal white adipose tissue (eWAT), and brown adipose tissue (BAT). (C) Fasting blood glucose (mmol/L), plasma triglycerides (mmol/L), total cholesterol (mmol/L), circulating lactic acid (mmol/L), and TNF-α (pg/mL) measured at sacrifice. (D) Histological quantification of adipocyte number per field in hematoxylin-eosin-stained eWAT sections and Liver steatosis scoring. (E) Blood glucose time course (mmol/L) following oral glucose challenge (2 g/kg; OGTT) and corresponding area under the curve (AUC, mmol/L×min). (F) Blood glucose time course during intraperitoneal insulin tolerance test (0.75 U/kg; IPITT) and corresponding AUC (mmol/L×min). Both tests were performed during the last week of the dietary intervention in CRL, HFHS, GSE1X, and GSE2X mice. Data are presented as mean ± SEM. Statistical comparisons by one-way ANOVA with Tukey’s post-hoc test unless otherwise stated. **p < 0.01, ****p < 0.0001. n = 8 mice per group.

HFHS induced a broad systemic metabolic syndrome profile (Fig. 1C). Fasting blood glucose was significantly increased in HFHS versus CRL mice (****p < 0.0001) while both GSE doses reduced significantly fasting blood glucose (****p < 0.0001 vs HFHS and **p < 0.0001 vs HFHS).

Plasma triglycerides were significantly reduced by GSE2X only (*p < 0.05 vs HFHS). Total cholesterol was increased by HFHS diet (****p < 0.001 vs CRL) and reduced by both GSE1X (**p < 0.01 vs HFHS) and GSE2X (*p < 0.05 vs CRL). Circulating lactic acid was significantly reduced by both GSE1X (***p < 0.001) and GSE2X (**p < 0.01) vs HFHS. Plasma TNF-α was increased in HFHS mice (*p < 0.05 vs CRL) and returned to CRL level in both GSE1X and GSE2X (*p < 0.05 each).

### GSE fully restores glucose tolerance and insulin sensitivity

To assess whole-body glucose homeostasis, oral glucose tolerance tests (OGTT) and intraperitoneal insulin tolerance tests (IPITT) were performed (Fig. 1E). HFHS mice exhibited severely impaired glucose tolerance, with area under the curve (AUC) values above those observed in the CRL group. Both GSE1X and GSE2X normalized OGTT-AUC (Fig. 1F), demonstrating a dose-equivalent restoration of glucose clearance capacity. Insulin sensitivity, assessed by IPITT-AUC, was severely compromised in HFHS mice, reflecting profound peripheral insulin resistance. Both GSE doses reversed this effect, GSE1X and GSE2X. Taken together, these results demonstrate that GSE exerts a powerful, coordinated restoration of homeostasis of the glucose–insulin axis.

### GSE remodels colonic inflammatory, antioxidant, tight junction, and enteroendocrine gene expression

Colonic mucosal gene expression revealed that the HFHS diet profoundly modified intestinal physiology at the transcriptional level (Fig. 2A). IL-6 expression was upregulated in HFHS (**p < 0.01 vs CRL), and GSE2X specifically suppressed this effect (**p < 0.01 vs HFHS). TNF-α was reduced in HFHS versus CRL (**p < 0.01), and in all the other groups. Colonic CAT gene expression was upregulated by HFHS (***p < 0.001 vs CRL), likely reflecting a compensatory response to oxidative stress. Both GSE doses normalized CAT (**p < 0.01 each vs. HFHS). Compared to CRL, a trend in the increase of the expression of SOD was observed in the HFHS group, while GSE2X specifically increased significantly the expression of SOD (*p < 0.001) compared to the CRL group. Tight junction proteins, which maintain the integrity of the paracellular intestinal barrier, were broadly altered by HFHS. Claudin expression was increased in the HFHS group (**p < 0.01 vs CRL) but not significantly reduced by either treatment. ZO-1 was markedly increased in the HFHS group (***p < 0.001 vs CRL) and normalized by GSE2X, but not by the lower dose. Occludin was increased only by HFHS and GSE2X. MUC2 was significantly increased by HFHS and decreased by the two doses of GSE, specifically GSE2X (***p < 0.001).

**Figure 2.**
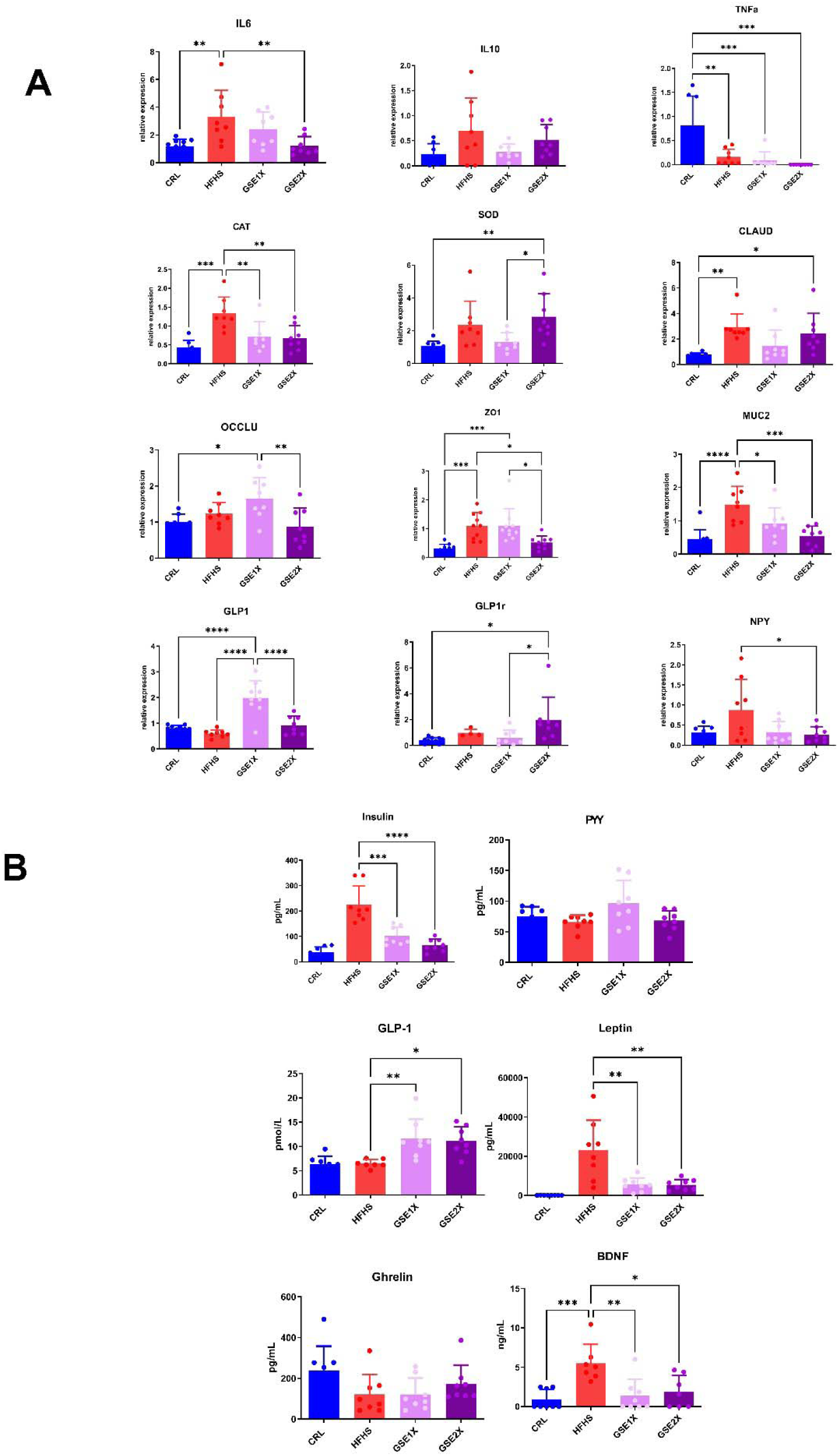
| GSE administration modulates colonic gene expression of inflammatory, oxidative stress, barrier integrity, enteroendocrine markers, and circulating hormones and neurotrophic factors. (A) Relative mRNA expression (normalised to housekeeping genes, RT-qPCR) in colonic tissue from CRL, HFHS, GSE1X, and GSE2X mice. Markers shown: IL-6, IL-10, TNF-α (inflammation); CAT, SOD (antioxidant defense); Claudin, Occludin, ZO-1 (tight junction integrity); MUC2 (mucus layer); GLP-1, GLP-1r (enteroendocrine signaling); NPY (neuropeptide). (B) Fasting plasma concentrations of insulin (pg/mL), peptide YY (PYY, pg/mL), glucagon-like peptide-1 (GLP-1, pmol/L), leptin (pg/mL), ghrelin (pg/mL), and brain-derived neurotrophic factor (BDNF, ng/mL) measured by ELISA or MSD multiplex electrochemiluminescence assay in CRL, HFHS, GSE1X, and GSE2X mice at sacrifice. Data are presented as mean ± SEM. Statistical comparisons by one-way ANOVA with Tukey’s post-hoc test unless otherwise stated. *p < 0.05, **p < 0.01, ***p < 0.001, ****p < 0.0001. n = 8 mice per group

Colonic GLP-1 mRNA expression was increased by GSE1X compared to all the other groups, (****p < 0.0001 vs CRL, HFHS, and GSE2X). This provides a direct mechanistic link between GSE1X supplementation, enhanced colonic enteroendocrine gene transcription, and elevated circulating GLP-1. GSE2X, by contrast, specifically upregulated GLP-1 receptor (GLP-1r) mRNA vs all other groups. NPY was reduced in the GSE2X compared to the HFHS group (*p < 0.05 vs HFHS),.

### GSE elevates circulating GLP-1, resolves hyperinsulinemia and hyperleptinemia, and normalizes plasma BDNF

Circulating hormone profiling revealed convergent endocrine responses to GSE treatment that mechanistically link adipose remodeling to gut–brain axis signaling (Fig. 2 B). Fasting hyperinsulinemia, a hallmark of peripheral insulin resistance in HFHS mice, was significantly reduced by GSE1X and GSE2, in concordance with the IPITT improvements observed in Fig. 1F. GLP-1 is an incretin hormone playing a central role in blood sugar regulation and appetite. Interestingly, GLP-1 was significantly increased by both GSE1X (**p < 0.01) and GSE2X(*p < 0.05) compared to the HFHS group, indicating enhanced GLP-1 signaling associated with GSE supplementation.

Leptin, an adipokine whose circulating concentration directly mirrors white adipose tissue mass, was significantly increased in HFHS mice compared to CRL (****p < 0.0001 vs HFHS). Both GSE doses markedly reduced hyperleptinemia induced by the HFHS diet (**p < 0.01 each). BDNF was increased in HFHS plasma compared to the CRL group (***p < 0.001 vs CRL) and was reduced significantly by both GSE doses compared to the HFHS group. PYY and ghrelin did not differ significantly across groups.

### GSE restores hepatic antioxidant defense and lipid-metabolic gene programs

Hepatic transcriptional profiling revealed that HFHS diet suppressed key antioxidant and catabolic gene expression while activating lipogenic pathways (Fig. 3 A). Hepatic CAT mRNA was decreased in the HFHS group (*p < 0.05 vs CRL) and was restored to the CRL level by both GSE doses. More strikingly, SOD expression was significantly decreased in the HFHS group (***p < 0.001 vs CRL), and both GSE doses induced a restoration of hepatic SOD, with GSE1X exceeding even CRL levels (**p < 0.01 vs CRL). Sterol regulatory element-binding proteins (SREBP) expression was decreased by HFHS and restored by GSE1X (*p < 0.05 vs HFHS) and GSE2X (**p < 0.01 vs HFHS) doses. Carbohydrate response element-binding protein (ChREBP) expression, which mediates glucose-driven lipogenesis, was significantly decreased in HFHS liver (***p < 0.001 vs CRL) and was fully restored to the CRL level by both GSE doses.

**Figure 3.**
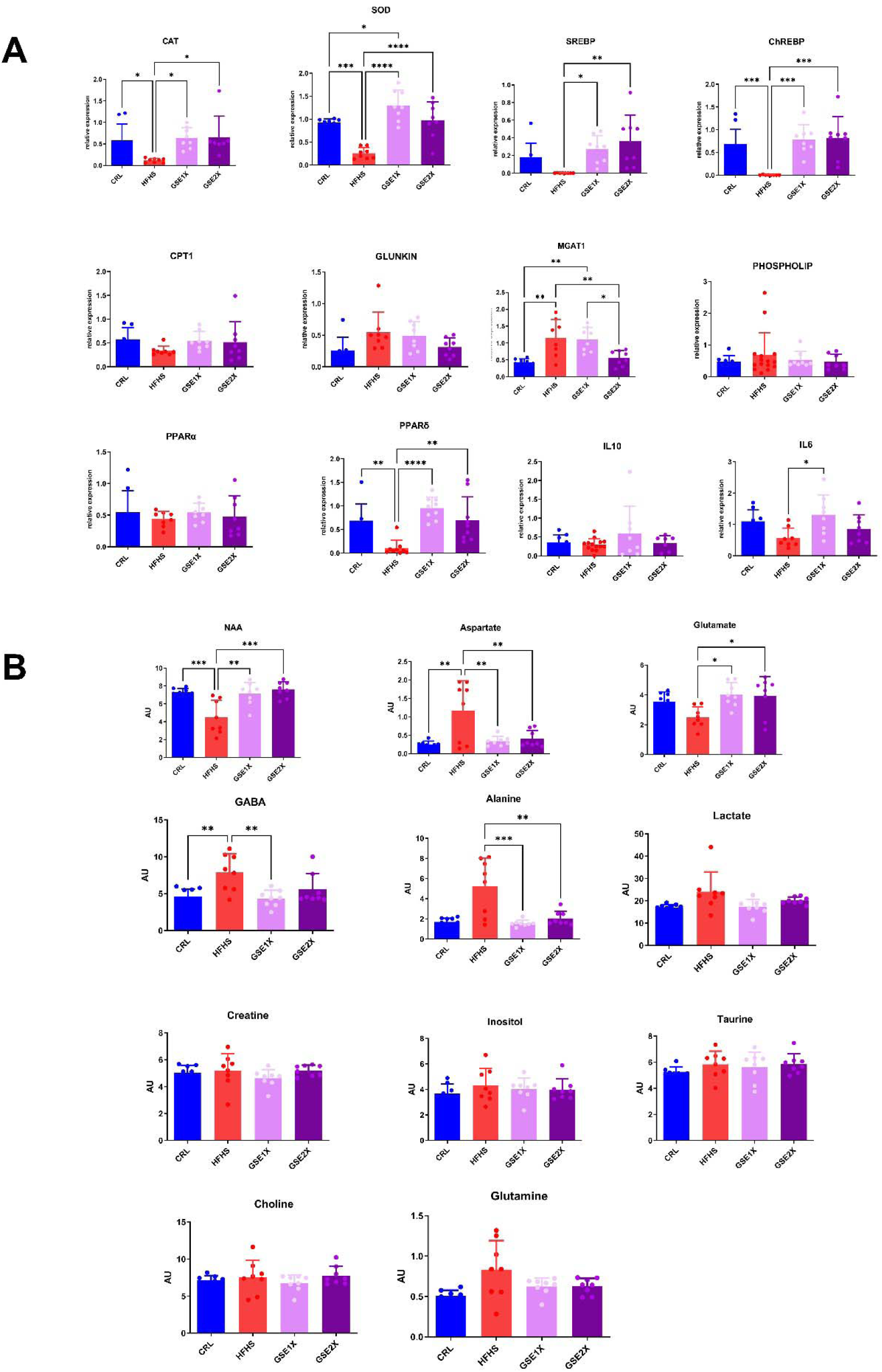
| GSE administration restores hepatic antioxidant defenses, lipid metabolism transcription, inflammatory signaling and reverses obesity-induced cerebral neurochemical dysfunction assessed by in vivo ¹H-MRS. (A) Relative mRNA expression (normalised to housekeeping genes, RT-qPCR) in liver tissue from CRL, HFHS, GSE1X, and GSE2X mice. Genes shown: CAT, SOD (antioxidant enzymes); SREBP, ChREBP (lipogenic transcription factors); CPT1, PPARα, PPARδ (fatty acid oxidation and nuclear receptors); MGAT1, Phospholipase (lipid metabolism); Glucokinase (glycolysis); IL-6, IL-10 (inflammatory cytokines). GSE supplementation reverses obesity-induced cerebral neurochemical dysfunction assessed by in vivo ¹H-MRS. Concentrations of eleven brain metabolites — N-acetylaspartate (NAA), aspartate, glutamate, GABA, alanine, lactate, creatine, inositol, taurine, choline, and glutamine — quantified by ex vivo ¹H High-Resolution Magic Angle Spinning (HR-MAS) NMR spectroscopy in CRL, HFHS, GSE1X, and GSE2X mice. Data are presented as mean ± SEM. Statistical comparisons by one-way ANOVA with Tukey’s post-hoc test unless otherwise stated. *p < 0.05, **p < 0.01, ***p < 0.001, ****p < 0.0001. n = 8 mice per group.

Monoacylglycerol acyltransferase 1 (MGAT1) was increased in HFHS compared to CRL group (**p < 0.01 vs CRL) and was specifically reduced by GSE2X (**p < 0.01 vs HFHS;*p < 0.05 vs GSE1X), mechanistically consistent with the GSE2X-specific reduction in plasma triglycerides observed in Fig. 1C. PPARδ was significantly downregulated in HFHS (**p < 0.01 vs CRL), and both GSE doses prevented this decrease induced by HFHS diet. Hepatic CPT1, GLUNKIN, PPARα, phospholipid synthesis, and IL-10 mRNA did not differ significantly across groups. A mild increase in IL-6 expression was detected in GSE1X-treated group (*p < 0.05 vs HFHS) but not in the GSE2X group.

### GSE restores brain metabolism

Quantification of ^1^H-NMR spectra from brain biopsies provided direct evidence that the HFHS diet impairs central neurometabolism and that GSE partially reverses these deficits (Fig. 3 B). NAA was significantly reduced in HFHS group (***p < 0.001 vs CRL) while GSE1X and GSE2X restored NAA to CRL levels. Aspartate was increased in HFHS compared to CRL (**p < 0.01 vs CRL) and was normalized by both GSE1X and GSE2X. Glutamate was reduced by HFHS, but this decrease was prevented by both GSE1X and GSE2X.

GABA was increased by HFHS (**p < 0.01 vs CRL), and GSE1X specifically normalized this effect, restoring the excitatory–inhibitory amino acid balance. Cerebral alanine tended to be increased by HFHS and was strongly reduced by both GSE1X (***p < 0.001 vs HFHS) and GSE2X (**p < 0.01 vs HFHS), returning to the level observed in the CRL group. Lactate, creatine, inositol, taurine, choline, and glutamine did not differ significantly across groups. These neurometabolic improvements, occurring in parallel with activation of the GLP-1 axis, normalization of leptin levels, and reduced plasma BDNF levels, as documented in Fig. 3, suggest an association between GSE-induced endocrine remodeling and improved central neurometabolic homeostasis.

### GSE administration profoundly remodels gut microbiota composition and diversity

Principal coordinates analysis (PCA) using Bray-Curtis dissimilarity confirmed complete separation of all four groups in ordination space (Axis 1 accounting for 43.48% and 46.53% of variance in two independent analyses, respectively), indicating that both diet and GSE treatments induced distinct microbial community states. Alpha diversity (observed richness) followed the order CRL > HFHS ≈ GSE1X >> GSE2X (****p < 0.0001 for all pairwise comparisons involving GSE2X vs all the compared groups; Fig. 4 A), indicating that GSE2X produced the most radical compositional reorganization of the microbiota. Inverse Simpson diversity, which weights evenness over richness, revealed a strikingly divergent pattern: GSE1X achieved the highest evenness of all groups (****p < 0.0001 vs both CRL and HFHS), while GSE2X showed the lowest (****p < 0.0001 vs both CRL and HFHS). GSE1X simultaneously maintains near-normal richness and achieves superior evenness, whereas GSE2X drives the most radical reduction in species number, demonstrating that the two doses induce distinct microbial restructuring effects rather than a simple dose–response continuum. Shannon, Simpson, and Bray diversity indices did not differ significantly across groups.

**Figure 4.**
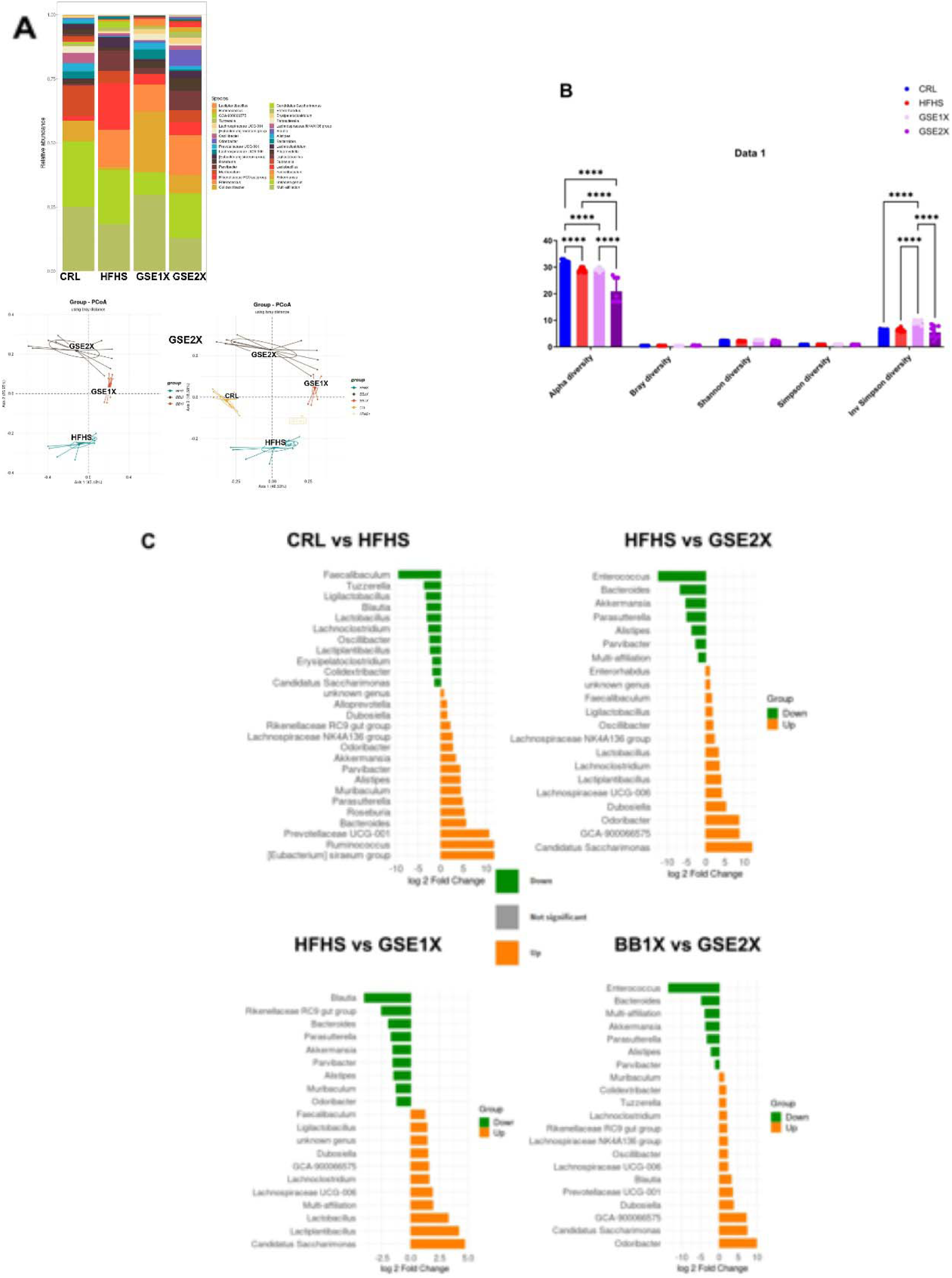
| GSE administration induces distinct gut microbiota community structures, diversity. and gut microbiota composition at the genus level. (A) Stacked bar charts showing relative abundance of dominant bacterial genera across CRL, HFHS, GSE2X, and GSE1X groups based on 16S rRNA V3–V4 region amplicon sequencing. Principal coordinate analysis (PCoA) plots based on Bray–Curtis dissimilarity; axis 1 explains 43.48% and 46.53% of total variance in the two analyses. Each point represents one animal; ellipses denote group dispersion. Beta diversity assessed by PERMANOVA. n = 8 mice per group.(B) Alpha diversity indices — observed species richness (Alpha diversity), Bray diversity, Shannon index, Simpson index, and inverse Simpson index — computed from 16S rRNA amplicon sequencing data of fecal samples from CRL, HFHS, GSE1X, and GSE2X mice. (C) DESeq2-based log_2_ fold change (log_2_FC) of differentially abundant genera for four pairwise comparisons: CRL vs. HFHS; HFHS vs. GSE2X; HFHS vs. GSE1X; GSE1X vs. GSE2X. Green bars (Down): genera more abundant in the first-named group; orange bars (Up): genera more abundant in the second-named group. Only genera reaching adjusted p < 0.05 (Benjamini–Hochberg correction) are displayed. 16S rRNA amplicon data, V3–V4 region, Illumina MiSeq. n = 8 mice per group

Differential abundance analysis at the genus level revealed a profound HFHS-induced dysbiosis relative to CTR animals (Fig. 4C): genera significantly enriched in HFHS included *Faecalibaculum*, *Tuzzerella*, *Ligilactobacillus*, *Blautia*, *Lactobacillus*, *Lachnoclostridium*, *Oscillibacter*, *Lactiplantibacillus*, *Erysipelatoclostridium*, and *Colidextribacter*, while genera significantly enriched in CTR included *Bacteroides*, *Akkermansia*, *Parasutterella*, *Alistipes*, *Prevotellaceae UCG-001*, *Ruminococcus*, and the *[Eubacterium] siraeum* group, reflecting a characteristic shift toward a metabolically unfavorable microbiome under HFHS feeding. Fig. 8 comparing HFHS vs GSE2X showed that genera i *Enterococcus*, *Bacteroides*, *Akkermansia*, *Parasutterella*, and *Alistipes* were markedly enriched in GSE2X compared to HFHS. While genera enriched in HFHS included *Lactobacillus*, *Lactiplantibacillus*, *Lachnoclostridium*, *Lachnospiraceae NK4A136 group*, *Dubosiella*, *Candidatus Saccharimonas*, *GCA-900066575*, and *Odoribacter*, confirming a robust restoration of health-associated commensal communities by GSE2X. Similarly, GSE1X treatment significantly enriched taxa, including *Blautia*, *Bacteroides*, *Akkermansia*, *Parasutterella*, *Alistipes*, and *Muribaculum* relative to (Fig. 4C) compared to the HFHS group. While genera significantly reduced in GSE1X included *Faecalibaculum*, *Ligilactobacillus*, *Lachnoclostridium*, *Lachnospiraceae UCG-006*, *Lactobacillus*, *Lactiplantibacillus*, and *Candidatus Saccharimonas*,. Direct comparison between GSE1X and GSE2X (Fig. 8) revealed that GSE2X-enriched genera included *Enterococcus*, *Bacteroides*, *Akkermansia*, *Parasutterella*, and *Alistipes*, while GSE1X-enriched genera included *Candidatus Saccharimonas*, *GCA-900066575*, *Dubosiella*, *Blautia*, *Prevotellaceae UCG-001*, and *Odoribacter*, highlighting a dose-dependent divergence in microbiota composition.

### Metabolomics reveals GSE bioavailability, microbial metabolite normalization, and restoration of mitochondrial acylcarnitine profiles

Untargeted metabolomic profiling revealed a clear disruption of the cecal metabolome in the HFHS group compared with control, as shown by the separation in the PCA scores plot (Fig. 5A) and the large number of significantly altered pathways (Fig S1A). Pathway enrichment analysis further indicated that HFHS feeding affected multiple metabolic pathways, including fatty acid-related pathways, steroid and bile acid metabolism, and amino acid metabolism (Fig. 5B). These changes were accompanied by marked shifts in representative metabolites spanning bile acids and steroids (Fig. 6A), lipid-derived species (Fig. 6B), and other secondary metabolites (Fig. 6D), consistent with a broad diet-induced remodeling of the cecal metabolic environment.

**Figure 5:**
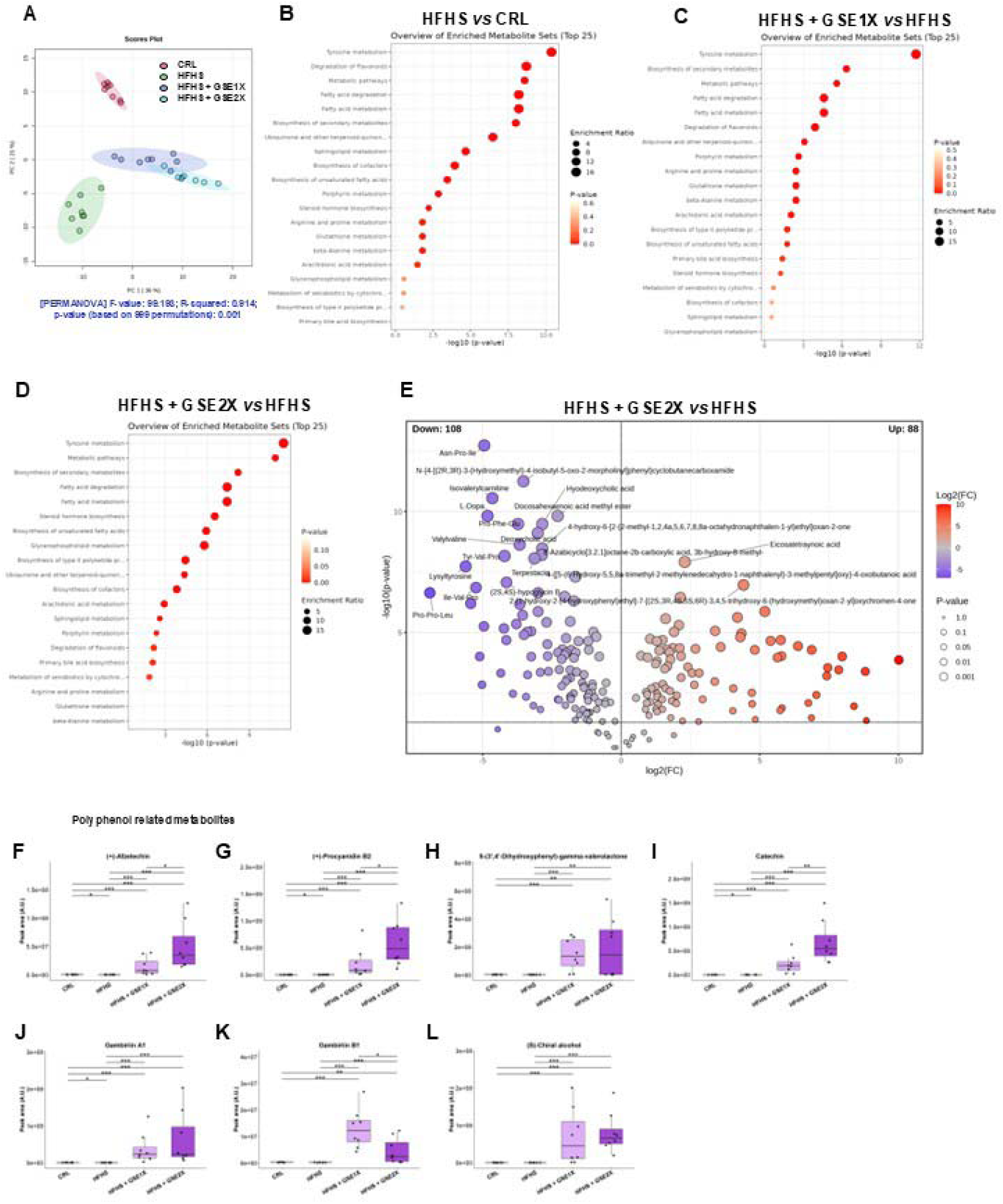
GSE administration reshapes the caecal metabolome of HFHS mice and modulates key metabolic pathways. (A) PCA score plot showing separation between CRL, HFHS and GSE-treated groups; ellipses indicate group dispersion and PERMANOVA statistics are shown in the panel.(B) Bubble plot of the 210 significant annotated features identified by Kruskal-Wallis testing across the four experimental groups, with significance represented by –log 10(p); the most significant metabolites are labeled. (C, E) Enrichment analysis of significantly altered metabolites, highlighting the top enriched metabolic pathways in CRL vs HFHS groups comparison (C) or HFHS vs GSE2X groups comparison (E). (D) Volcano plot of differential metabolites between HFHS + BB2X vs HFHS (p-value threshold: 0.05).

**Figure 6:**
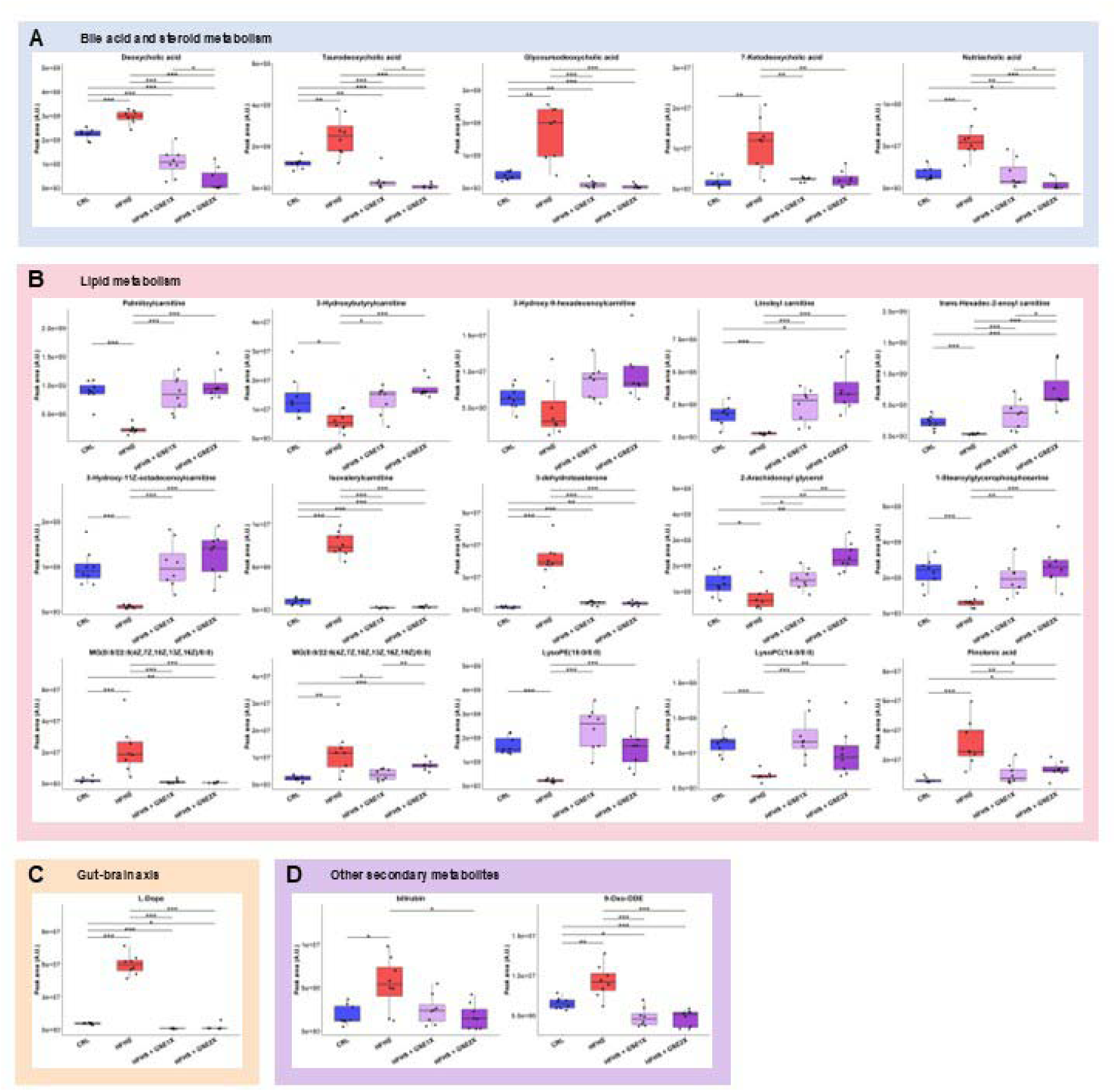
Representative annotated metabolites selected from the significant features. (A-X) Boxplots of discriminant metabolites show marked differences between CRL and HFHS, and between HFHS and HFHS+ BB1X/BB2X mice across lipid, steroid, bile acid, and redox-related pathways, indicating that grape seed extract treatment substantially alters the HFHS-induced metabolic profile. Annotated metabolites shown were selected to represent the main metabolic classes altered by the HFHS diet and modulated by GSE treatment, in order to highlight the most biologically relevant trends. Overall differences were analyzed using the Kruskal–Wallis test, followed by pairwise Wilcoxon rank-sum (Mann–Whitney) tests with Benjamini–Hochberg correction for multiple comparisons. Significance levels are indicated as P < 0.05 (), P < 0.01 (), P < 0.001 (), and P < 0.0001 (****).

Compared with HFHS, GSE treatments partially shifted the metabolomic profile toward the control group, with a more pronounced effect for GSE2X than for GSE1X (Fig. 6A). Hierarchical clustering of the 210 annotated discriminant metabolites confirmed the global metabolomic changes observed in the HFHS model and further suggested a stronger effect of GSE2X than GSE1X (Fig. S1B). Notably, the HFHS samples appeared closer to the control group than to the GSE-treated groups, indicating that GSE treatments drove a distinct metabolic shift rather than a complete normalization toward the control profile (Fig. 1B). The subsequent analyses focused on the HFHS versus HFHS + GSE2X comparison, which showed 108 down– and 88 up-regulated metabolites when compared to HFHS (Fig. 5E), and a partial normalization of several altered metabolic pathways including fatty acid metabolism, bile acid and amino acid synthesis, and steroid and lipids biosynthesis (Fig. 5D). This recovery was also reflected in the comparative enrichment analysis between HFHS and HFHS + GSE2X, which suggested attenuation of the metabolic perturbations induced by the HFHS diet. Overall, these data indicate that the HFHS diet strongly perturbs the cecal metabolome, while both GSE doses exert a partial corrective effect on the metabolic signature.

Dose-dependent absorption of GSE polyphenols was confirmed by significantly elevated polyphenol related metabolites, including afzelechin (Fig.5F), procyanidin B2 (Fig5G), 5-(3’,4’-Dihydroxyphenyl)-gamma-valerolactone (Fig. 5H), catechin (fig5I), gambiriin A1 (Fig. 5J) and B1 (Fig. 5K) and chiral alcohol (Fig. 5L) in both GSE groups vs HFHS, with GSE2X predominating for most analytesNotably, gambirin B1 showed an inverse pattern, accumulating more in GSE1X than GSE2X, suggesting differential gut microbial biotransformation at the two doses. 5-(3’,4’-Dihydroxyphenyl)-gamma-valerolactone was equivalently elevated in both GSE groups versus CRL and HFHS. These results support a dose-dependent absorption and microbial metabolism of GSE polyphenols, and we next examined the broader impact of treatment on other metabolite classes.

Regarding other microbiota-derived metabolites, the significant features highlighted coordinated changes in bile acid and steroid metabolism, lipid metabolism, and gut-brain axis-associated compounds following GSE2X treatment in HFHS-fed mice (Fig. 5D and 6). In the bile acid and steroid metabolism panels, HFHS strongly increased deoxycholic acid, taurodeoxycholic acid, glycoursodeoxycholic acid, 7-ketodeoxycholic acid, and pinolenic acid, whereas GSE partially counteracted these changes, particularly in the GSE2X group (Fig. 6A). Similarly, lipid-related metabolites such as palmitoylcarnitine, 3-hydroxybutyrylcarnitine, 3-hydroxy-9-hexadecenoylcarnitine, linoleyl carnitine, trans-hexadec-2-enoyl carnitine, 2-arachidonoyl glycerol, and several lysolipids showed reduced levels with the HFHS diet that were partially normalized by GSE, again with the strongest effect generally observed for GSE2X (Fig. 6B). Isovalerylcarnitine, 3-dehydroteasterone, and monoacylglycerols (MG) levels were increased in the HFHS diet group, while both GSE groups seemed to partly suppress this effect (Fig. 6B). Finally, gut-brain axis-associated metabolite L-DOPA showed a significant increase in the HFHS group when compared to the control, which was restored by GSE, suggesting that the polyphenol extract may also influence metabolic pathways relevant to host-microbiota communication. Among the other secondary metabolites, bilirubin, 5-oxo-delta-bilirubin, and 9-Oxo-ODE also differed significantly across groups, supporting additional modulation of oxidative and secondary metabolic pathways in HFHS animals, which were partly restored by GSE treatments (Fig. 6D).

## Discussion

GSE at the two doses reduced diet-induced body weight gain in a strictly dose-dependent manner. The OPC dose effect aligns with prior reports of interventions in high-fat diet models ^35,36^. In our study of physiological parameters and weight gain, the GSE2X dose is clearly more effective and closer to the CRL group. BAT normalization, however, occurred exclusively at the higher dose, consistent with the threshold dependency of thermogenic activation. The observed dose-response pattern can be explained by the parahormetic mode of action of oligomeric procyanidins, whereby increasing luminal doses recruit a wider array of redox-sensitive and receptor-dependent signaling pathways ^36,37^. The present data confirm that dose escalation is associated with enhanced biological responses across multiple metabolic and microbial endpoints. The dominance of OPCs and OPC metabolites in the ceca of GSE-treated mice was demonstrated. The dominance of these metabolites at the GSE2X dose confirms greater intestinal exposure to OPC-derived metabolites, which may contribute to the stronger microbiota and host responses observed at this dose.

In our model, in the blood, the decreases in TNF-α with the two GSE doses were correlated with decreases in glucose, triglycerides, and cholesterol. In contrast, lactatemia was significantly increased by HFHS and decreased by both treatments. TNF-α disrupts metabolism by inhibiting carbohydrate metabolism, lipogenesis, adipogenesis, and thermogenesis, while stimulating lipolysis and impairing the endocrine functions of adipose tissue, thereby compromising its ability to store excess energy ^38^.

This proinflammatory state in the HFHS group, characterized by high TNF-α levels, can be either the cause or the consequence of insulin resistance ^5–7,39^. In our study, the high insulin levels in the HFHS group were markedly reduced by both GSE treatments. This improvement in insulin resistance may be a milestone in maintaining glucose homeostasis. The normalization of lactatemia further indicates an improvement in systemic metabolic homeostasis, potentially reflecting enhanced oxidative metabolism ^40^.

Circulating cholesterol levels were reduced with both doses, whereas triglyceride levels were reduced only in the GSE2X group. This dissociation maps directly onto a dose-specific hepatic mechanism: MGAT1, a rate-limiting enzyme in intestinal and hepatic triglyceride re-esterification, was reduced only by GSE2X. The liver exhibited a coordinated antioxidant and transcriptional response to GSE treatment. Hepatic superoxide dismutase (SOD) activity was elevated (p<0.0001 versus HFHS), exceeding even control levels and indicating a hormetic induction of the antioxidant defense program, a mechanism characteristic of procyanidin-class polyphenols ^37,41^. Catalase was also normalized. PPARδ, a nuclear receptor governing mitochondrial fatty acid β-oxidation ^42^, was restored by GSE1X (p < 0.0001), providing a transcriptional basis for improved lipid catabolism.

ChREBP, a carbohydrate-responsive element-binding protein that drives de novo lipogenesis in response to excess fructose and glucose flux ^43^ was normalized at the mRNA level in the liver, indicating reduced lipogenic pressure. MGAT1 suppression by GSE2X accounts for the selective reduction in plasma triglycerides. SREBP elevation in the GSE2X group could reflect an adaptive transcriptional response to altered hepatic lipid handing, as SREBP-mediated gene programs are subject to feedback regulation by the sterol content of the endoplasmic reticulum membrane ^44^. Polyphenols have been previously shown to attenuate hepatic steatosis in diet-induced models through overlapping mechanisms ^45^.

The HFHS diet induced a profound remodeling of the cecal metabolome, consistent with the broad metabolic and inflammatory phenotype observed in the animals. This effect was not restricted to a few individual metabolites, but rather involved coordinated changes in bile acid and steroid metabolism, lipid handling, and gut–brain axis-related compounds, indicating a systemic disturbance of intestinal biochemical homeostasis. Among the most affected pathways, bile acid metabolism was strongly altered, with changes in deoxycholic acid, taurodeoxycholic acid, glycoursodeoxycholic acid, and 7-ketodeoxycholic acid ^46^. Because bile acids are not only digestive molecules but also endocrine-like regulators of glucose and lipid metabolism, their dysregulation may contribute to the metabolic impairment induced by HFHS feeding ^47^. Similarly, the increase in lipid-related metabolites such as acylcarnitines and monoacylglycerols suggests altered fatty acid utilization and incomplete lipid catabolism, which are hallmarks of metabolic dysfunction. The colon represented a primary locus of anti-inflammatory activity ^48^. In this study, GSE-reduced colonic IL-6 expression, supporting a local anti-inflammatory effect of the extract. The blood, colon, and liver parameters collectively reflect a broad improvement of metabolic and inflammatory homeostasis.

Elevated circulating GLP-1 was observed in both treatment groups, suggesting a potential contribution of enteroendocrine signaling to the observed metabolic improvements. GLP-1 is a potent insulinotropic hormone secreted by intestinal L-cells in response to luminal nutrients and microbial signals ^49^.

In a previous study, we demonstrated the positive effects of the GSE in a normal mouse model on the intestinal GLP1 expression ^25^. GLP-1 secretion in obese people is severely attenuated compared to lean persons ^50^. *Akkermansia muciniphila* cell-free supernatant (CFS), particularly its P9 protein, markedly enhances GLP-1 secretion in NCI-H716 cells and primary human intestinal epithelial cells, and increases Gcg and Pcsk1 expression in the ileum of high-fat diet–fed mice ^51^.

In our study, *Akkermansia muciniphila* increased significantly and in a clear dose-dependent manner, which may account for the concomitant rise in circulating GLP-1. These results are consistent with the recent clinical trial by Suenaert et al., which reported improvements in insulin sensitivity, body composition, and GLP-1 production in patients with metabolic syndrome receiving pasteurized *A. muciniphila* ^52^. In this study, intestinal and circulating GLP-1 levels were significantly improved following GSE supplementation. GGSE2X induced colonic GLP-1r expression, suggesting enhanced sensitivity of target tissues to circulating ligand rather than increased hormone biosynthesis. Both strategies are consistent with the emerging understanding that dietary polyphenols can modulate gut hormone cascades through microbiota-dependent mechanisms ^53,54^.

BDNF in plasma was elevated in HFHS animals, which may reflect an adaptive response to metabolic stress, which is a non-obvious effect. Both GSE doses normalized BDNF level, indicating normalization of this altered neurotrophic profile ^55^. Leptin normalization observed in GSE-treated animals may reverse energy homeostasis. GSE normalizes gut-derived L-DOPA levels, highlighting a marked effect on microbiota-associated catecholamine precursor metabolism ^56,57^.

At the microbiota level, GSE1X maintained high species richness and achieved the highest diversity score, indicating a diverse and evenly distributed microbiota, whereas GSE2X induced a low-richness, uneven community dominated by a few taxa. Despite this reduced diversity, GSE2X still conferred strong metabolic benefits, showing that global diversity metrics alone do not fully explain physiological effects. The low alpha-diversity in GSE2X can be explained by two converging mechanistic forces. First, GSE exerts documented antimicrobial activity against a range of taxa ^25^Second, through the marked expansion of *Akkermansia muciniphilia (A. muciniphila),* the keystone of microbial anti-obesity ^51,58,59^. Even in humans, supplementation with *A. muciniphila* improved insulin sensitivity, reduced insulinemia and plasma total cholesterol ^60^, and helped maintain weight loss in people with overweight and obesity at the beginning of the study ^61^. Many studies have demonstrated an intestinal bloom of *A. muciniphila* induced by various OPCs and polyphenols ^62–64^. In the present study, the dose-dependent effect of GSE on the intestinal bloom of *A. muciniphila* was particularly striking, with the highest dose inducing the most pronounced expansion of this mucus-associated commensal bacterium. In a previous study using the same dose of GSE in lean animals fed with a chow diet, we didn’t see any increase in *A. muciniphila*.

A recent study demonstrated that *A. muciniphila* uses dietary polyphenols as xenosiderophores to enhance iron uptake ^65^. What could explain the different effect of *A. muciniphila*, which increases in animals fed an HFHS diet but not in lean animals. Indeed, the HFHS model increases iron bioavailability at the intestinal level ^66,67^. In our study, we demonstrated that the more OPC we administer, the more it is detected in the intestine by metabolomic analysis, and the more it modulates the gut microbiota, with an expansion of *A. muciniphila*. The abundant presence of iron and bioavailable OPCs may contribute to the expansion of *A. muciniphila* in HFHS models.

Many studies have demonstrated the important role of *A. muciniphila* in the gut-brain axis ^68,69^. In our study, the expansion of *A. muciniphila* occurred concomitantly with marked improvements in several neurochemical parameters.

HR-MAS provided direct neurochemical evidence that dietary obesity is not merely a peripheral metabolic disorder. Twelve weeks of HFHS feeding produced measurable alterations in brain tissue metabolites, including a significant reduction in NAA. NAA is synthesized exclusively in neuronal mitochondria and exported to oligodendrocytes, where it serves as an acetyl group donor for myelin synthesis ^70^. Its concentration in brain tissue is therefore a dual readout of neuronal mitochondrial function and axonal integrity. NAA reduction in the HFHS brain indicates both energetic failure at the neuronal level and potential compromise of myelination ^71^. GSE2X restored brain NAA to control levels, suggesting a beneficial effect on neuronal metabolic integrity.

To our knowledge, this is the first study reporting a positive effect of GSE in the HFHS model on the central nervous system axis. It is also known that the HFHS diet further disrupted the excitatory/inhibitory neurotransmitter balance in the brain. Aspartate, an excitatory amino acid and NMDA receptor co-agonist, was elevated in HFHS brains, raising the risk of excitotoxic signaling. Simultaneously, GABA was elevated and glutamate was reduced, a pattern indicative of a dysfunctional compensatory shift in the excitation/inhibition balance rather than a physiological predominance of inhibition. Both GSE doses normalized aspartate and alanine. GSE1X specifically restored GABA to control levels, while GSE2X restored glutamate and normalized NAA, suggesting that the two doses engage complementary neurotransmitter normalization programs. Plasma BDNF was paradoxically elevated in HFHS animals, consistent with a neuroinflammatory compensatory response in which BDNF is acutely upregulated to sustain neuronal survival under metabolic stress. Both GSE doses normalized BDNF, indicating resolution of the underlying neuroinflammatory (involved in the development in several brain disorders) drive rather than suppression of a beneficial signal. These findings confirm that obesity-associated neurochemical alterations occur in parallel with major changes in gut physiology and microbiota composition, and that both can be improved by GSE supplementation, as conceptualized in the comprehensive framework of Cryan *et al.* ^55^.

Multiple convergent pathways likely link the intestinal effects of GSE to brain metabolism. Beyond their role in glucose homeostasis (Holst, 2007), GLP-1 receptors are widely expressed in the brain^49^. Recently, it has been demonstrated that GLP-1 receptor knockout mice exhibit impaired cerebral glucose metabolism, demonstrating that intact GLP-1 signaling is required to maintain normal brain bioenergetics ^72^. Restoration of peripheral GLP-1 signaling by GSE could therefore help normalize cerebral metabolic pathways, ultimately improving neuronal mitochondrial metabolism. Such metabolic restoration is consistent with the recovery of metabolic profiles, in particular glutamate and GABA levels observed in this study, and with the increase in N-acetylaspartate (NAA), a marker of neuronal mitochondrial integrity and function. Moreover, Isovalerylcarnitine, a short-chain acylcarnitine derived from leucine catabolism (a branched-chain amino acid – BCAA), was markedly elevated in HFHS mice and normalized by both GSE treatment, suggesting restoration of leucine catabolism. This finding may be particularly relevant because leucine is a major precursor of cerebral glutamate and GABA through branched-chain aminotransferase-mediated transamination. Consistent with this metabolic link, we previously demonstrated that alterations in peripheral BCAA metabolism modulate cerebral GABA metabolism ^73^. Although leucine itself was not quantified in the present study, normalization of isovalerylcarnitine may therefore reflect improved BCAA metabolism that contributes to the restoration of glutamate and GABA homeostasis. Given that glutamate and GABA are the principal excitatory and inhibitory neurotransmitters in the mammalian brain, respectively, maintaining a proper balance between these neurotransmitter systems is essential for normal neuronal network activity. Disruption of the glutamate–GABA equilibrium has been implicated in cognitive dysfunction, anxiety-like behavior, and other neurobehavioral alterations ^74^.

The present findings indicate that GSE supplementation counteracted many of these HFHS-induced alterations in a dose-dependent manner, with GSE2X generally producing a stronger and more coherent effect than GSE1X. This is fully consistent with the physiological data, in which the higher dose was more effective at reducing body weight gain, normalizing circulating metabolic markers, and improving tissue-level readouts such as BAT function, inflammation, and endocrine parameters. At the metabolomic level, GSE partially restored the cecal profile toward that of control animals, suggesting that its beneficial effects are at least partly mediated by correction of the intestinal metabolic environment. In particular, the partial suppression of HFHS-induced increases in isovalerylcarnitine, 3-dehydrotestosterone, and monoacylglycerols supports normalization of lipid– and steroid-related metabolism, while the broader normalization of bile acid-related compounds is consistent with improved metabolic signaling and host– microbiota crosstalk.

## Conclusion

This study demonstrates that oral supplementation with GSE, an oligomeric procyanidin-enriched extract, exerts potent, dose-dependent, and mechanistically distinct effects against diet-induced obesity in C57BL/6J male mice fed a high-fat, high-sugar diet. At both doses, GSE significantly attenuated body weight gain, visceral adiposity, and metabolic syndrome parameters, including fasting hyperglycemia, dyslipidemia, hyperinsulinemia, and systemic inflammation, while restoring glucose tolerance and insulin sensitivity to near-control levels.

At the lower dose, the predominant mechanism involves the selective enrichment of a diverse and evenly distributed gut microbial community, characterized by the expansion of health-promoting taxa, including *A. muciniphila*, *Bacteroides*, and *Alistipes*, coupled with transcriptional induction of colonic and plasmatic GLP-1 production and the dose-specific modulation of bile acid profiles toward cytoprotective secondary bile acids. At the higher dose, a more radical restructuring of the microbiota, dominated by *A. muciniphila* expansion converges with induction of hepatic antioxidant defenses (SOD), suppression of MGAT1-dependent triglyceride re-esterification, enhanced colonic GLP-1 receptor expression, and full restoration of neuronal N-acetylaspartate, a direct marker of cerebral mitochondrial integrity and axonal viability. Both doses converge on shared outcomes: normalization of the enteroendocrine GLP-1 axis, resolution of hyperleptinemia, suppression of microbial L-DOPA overproduction, and restoration of acylcarnitine profiles indicative of recovered mitochondrial fatty acid β-oxidation.

Critically, this study provides the first direct evidence that a gut-acting polyphenol compound can reverse obesity-induced neurochemical dysfunction in the brain, as demonstrated by *ex vivo* ¹H HR-MAS NMR spectroscopy. These findings establish a comprehensive, multi-axis model of action for this specific GSE that simultaneously engages the gut microbiota, the gut-liver axis, and the gut-brain axis through mechanistically integrated and dose-tunable pathways.

Taken together, these preclinical results position this GSE as a promising prebiotic candidate for the dietary management of obesity and its metabolic and neurological comorbidities. Clinical trials enrolling overweight and obese populations, incorporating both metabolic and neurocognitive outcomes, are warranted to determine whether these coordinated effects translate to benefit in humans.

## Materials and methods

BERKEM kindly offered GSE powder. The composition of this OPCs-rich GSE was previously described ^24^.

### Animals

Animal experiments were carried out in accordance with the rural and maritime fishing codes R.214-87 to R.214-126 25/04/2025 – Experimental Animal Ethics Committee of Bordeaux University, APAFIS #50103.

Five-week-old C57Bl/6 male mice (Janvier, Lorient, France) were housed in individually ventilated cages (IVCs) under specific pathogen-free (SPF) conditions, with ad libitum access to food and water, maintained at a controlled temperature (22 ± 2°C) under a 12-h light/dark cycle.

After two weeks acclimation period (during which animals were fed with chow diet A04), mice were randomly assigned to 4 experimental groups (n = 8, 4 mice/cage):

– Chow diet group **CRL**: was fed under chow diet A04 (SAFE, AUGY, France) and daily gavaged with skim milk as a vehicle.
– The high-fat high-sucrose group **HFHS** was fed the HFHS diet 292H (SAFE, AUGY, France) 46% lipids, 38.5% carbohydrates, and 15.5% proteins; (4.4 kcal/g) (whole composition in Supplemental Material) and daily gavaged with skim milk as a vehicle.
– The high-fat, high-sucrose **GSE1X** group: HFHS treated daily with 1 g/kg/day GSE (1X dose) by gavage in skim milk.
– The high-fat, high-sucrose **GSE2X** group: HFHS treated with 2 g/kg/day GSE (2X dose).

Body weight and food intake were measured three times weekly. Feces were collected just before the first gavage (t = 0), and then after 1, 4, and 8 weeks of treatment. At the end of the 12-week gavage protocol, mice were deeply anesthetized with isoflurane (5% in oxygen) and euthanized by cervical dislocation. Blood was collected by cardiac puncture, in EDTA-coated tubes; tubes were centrifuged, and plasma fractions were stored at –80°C. At the time of the sacrifice, a comprehensive panel of tissues was harvested, including the liver, brain, white adipose tissue (WAT) depots (epididymal, inguinal, mesenteric, and retroperitoneal), brown adipose tissue (BAT), and caecum. All collected tissues were immediately snap-frozen in liquid nitrogen and stored at –80°C until further analysis. Selected tissue portions were preserved under specific conditions according to downstream applications: samples designated for gene expression analysis were immersed in RNAlater® stabilization solution (liver, colon, and ileum), while those intended for histological examination were fixed in 4% paraformaldehyde (BAT, liver, colon, and ileum).

### Physiological analysis

Oral glucose tolerance test (OGTT) was performed 7 days before the end of the protocol. Mice were fasted for 12 h (9 pm – 9 am), and the glucose tolerance test was initiated by gavage with 1 g glucose/kg body weight. Glycemia was measured, and blood samples were collected at each time point (t = 0, 15, 30, 60, 90 min) ^75^.

Intraperitoneal Insulin Tolerance Test (IPITT) was performed at week 10 of dietary intervention, as previously described ^75^. Briefly, mice were fasted for 6 h (8 am – 2 pm) in clean cages with free access to water. Body weight was recorded immediately before the test to calculate the individual insulin dose. A working insulin solution was freshly prepared on the day of the experiment by diluting human insulin (Humulin® R, 100 U/mL; Eli Lilly) in sterile 0.9% (w/v) NaCl saline to a final concentration of 0.1 U/mL. Fasting blood glucose was measured at baseline (t = 0 min) from a small incision at the femoral vein using a calibrated glucometer (Accu-Chek® Aviva, Roche Diagnostics) after discarding the first blood drop. Each mouse then received an intraperitoneal injection of insulin at a dose of 0.75 U/kg body weight for lean mice and 1.0 – 1.5 U/kg for HFD-fed obese mice, in a volume of 10 µL/g body weight, using a 28-gauge insulin syringe. Blood glucose was subsequently measured at t = 15, 30, 60, and 90 min post-injection by gently removing a small blood drop from the femoral vein and collecting (<5 µL) on a new glucometer test strip. Animals exhibiting blood glucose below 20 mg/dL or signs of severe hypoglycemia (trembling, loss of consciousness) were immediately rescued by intraperitoneal injection of 300 µL of 20% (w/v) glucose solution and excluded from subsequent analyses. Food was reintroduced into the cages upon completion of the test, and animals were monitored for at least 2 h post-procedure ^75^.

### Histological analysis

Liver and epididymal white adipose tissue (eWAT) were collected at sacrifice and immediately fixed in 4% formaldehyde for 24 h at 4°C. Following three washes in phosphate-buffered saline (PBS), tissues were stored in 70% ethanol at 4°C until paraffin embedding. Tissue sections of 4 µm thickness were cut and stained with hematoxylin-eosin-saffron (HES) for histomorphological analysis. Hepatic steatosis was graded according to the NAFLD Activity Score (NAS) system as described by Kleiner et al ^76^, with steatosis scored on a scale of 0 to 3 corresponding to < 5%, 5 – 33%, 33 – 66%, and > 66% of hepatocytes containing lipid vacuoles, respectively. All liver sections were evaluated under light microscopy at × 10 and × 20 magnification by an independent observer blinded to experimental group assignment. For eWAT, adipocyte hypertrophy was assessed by measuring the mean cross-sectional cell area on HES-stained sections using image analysis software. Crown-like structures (CLS) were quantified. All histological scoring and morphometric analyses were performed blindly by an independent observer^62^.

### ELISA analysis

Circulating concentrations of metabolic hormones, gut peptides, neurotrophic factors, and inflammatory cytokines were simultaneously quantified using the Meso Scale Discovery (MSD) electrochemiluminescence (ECL) immunoassay platform (Meso Scale Diagnostics, Rockville, MD, USA). For the quantification of labile peptides susceptible to rapid *ex vivo* proteolysis specifically GLP-1 and active ghrelin blood was drawn into pre-chilled EDTA tubes supplemented with DPP-IV inhibitor (10 µL/mL blood; Merck Millipore) and phenylmethylsulfonyl fluoride (PMSF, 1 mM final concentration) immediately upon collection; all remaining analytes (total GLP-1, PYY, leptin, insulin) were collected in standard pre-chilled EDTA tubes, while serum was recovered in separator tubes for cytokine analyses (IL-6, TNF-α, BDNF). Following centrifugation at 1,300 × g for 15 min at 4°C, plasma and serum aliquots were snap-frozen in liquid nitrogen and stored at −80°C until analysis. All assays were performed according to the manufacturer’s recommendations using the following kits: V-PLEX GLP-1 Active (ver. 2), MSD Total GLP-1 (ver. 2), Active Ghrelin, PYY, Mouse Metabolic Panel (Leptin/Insulin, K15124C), V-PLEX Pro-Inflammatory Panel 1 Mouse (IL-6 and TNF-α, K15048D), and BDNF Singleplex (Meso Scale Diagnostics). Briefly, MSD plates pre-coated with capture antibodies were blocked with MSD Blocker A solution for 30 min, then washed 3 times with PBS-Tween 0.05% (PBST). Samples were loaded in duplicate at appropriate dilutions alongside a 7-point, 4-fold serial-dilution standard curve and incubated for 2 h at room temperature on an orbital shaker (700 rpm) in the dark. After three PBST washes, SULFO-TAG-conjugated detection antibodies were added and incubated for an additional 2 h under identical conditions. Following a final wash cycle, 150 µL/well of 2× MSD Read Buffer T was dispensed, and plates were immediately analyzed on a MESO QuickPlex SQ 120 reader using Discovery Workbench v4.0 software (Meso Scale Diagnostics). Analyte concentrations were calculated using 4-parameter logistic (4PL) regression of the standard curve; runs with intra– or inter-assay coefficients of variation exceeding 15% were excluded. Results are expressed as pg/mL or ng/mL of plasma or serum.

### Biochemical analyses

Biochemical analyses were performed by the metabolomics and clinical biochemistry platform (University of Bordeaux, France) using standardized automated colorimetric and enzymatic assays on a clinical biochemistry analyzer. Free fatty acids (FFA/AGL) were quantified using an enzymatic colorimetric kit based on the acyl-CoA synthetase/acyl-CoA oxidase reaction (NEFA-C kit, Wako Diagnostics), with results expressed in mmol/L. Total cholesterol and LDL-cholesterol concentrations were measured by direct enzymatic colorimetric methods, with LDL-cholesterol calculated using the Friedewald equation when triglyceride levels were below 3.9 mmol/L [C-LDL = CT − C-HDL − (TG/2.2)]. Fasting plasma glucose was determined by the hexokinase/glucose-6-phosphate dehydrogenase (G6PDH) enzymatic method. Triglycerides were assayed using a glycerol-phosphate oxidase (GPO-PAP) enzymatic method. Plasma lactate (lactatemia) was measured by an enzymatic L-lactate dehydrogenase (L-LDH) method, with results expressed in mmol/L. Fructosamine, a marker of medium-term glycemic control reflecting mean plasma glucose over the preceding 2–3 weeks, was quantified by the nitroblue tetrazolium (NBT) reduction method in alkaline conditions, using a polyglycated lysine calibrator, with results expressed in µmol/L. All assays were performed in duplicate, and results are expressed as mean ± SEM per experimental group.

### RNA extraction

Tissue samples (liver, colon, ileum), stored in RNA later at –20°C, were thawed on ice before being transferred into 1 mL of Trizol containing 1.4 mm ceramic beads (Lysing matrix D, FastPrep, MP). For hepatic tissue, frozen samples were directly added to Trizol to avoid RNA degradation. Tissue disruption was achieved by mechanical action using a bead beater (Ozyme France). After adding chloroform, the RNA-containing upper phase was transferred to a tube containing isopropanol. Total extracted RNA was then isolated with RNeasy Minikit QIAGEN, according to the manufacturer’s instructions. RNA samples were finally treated with DNase (Turbo DNA-free, Ambion, Inc.) and converted to cDNA from 1 µg of RNA using the SuperScript IV reverse transcriptase system (Thermo Scientific, Vilnius, Lithuania) ^25^.

### qPCR analysis

The reaction mixture comprised iTaqTM Universal SYBR® Green Supermix (Bio-Rad, France), forward and reverse primers (200 nM each), and q.s. RNase/DNase-free water (Sigma) to a final volume of 12 µL. Quantitative real-time PCR (qPCR) reactions were performed using a CFX96TM Touch System (Bio-Rad), and the amplification program was 95°C for 3 min; 40 cycles of 95°C for 30 s, 55-60-61 or 62°C for 20 s, and 72°C for 10 s; and melting curves were obtained at the end with reads performed every 0.5°C (hold 2 s) from 70 to 90°C. To analyze gene expression, 1 µL of cDNA was added per well, and target gene copy numbers were normalized to housekeeping genes.

### DNA extraction

Around 100 mg of feces were accurately weighed and homogenized in Tris-EDTA buffer (Tris 0.1 mM, pH 8; EDTA 1M (Sigma); 1 mL of buffer for 200 mg of feces). Lysozyme (1:100, 300 mg/mL (Sigma)) was added to the mixture, which was incubated at 37°C for 1 h. Then, 200 µL of the mix was added to the NucleoSpin Soil Bead tube, and DNA was isolated using the NucleoSpin® Soil kit (Macherey-Nagel) according to the manufacturer’s instructions ^77^.

### DNA sequencing

Bacterial community profiling was carried out by targeting the hypervariable V3– V4 region of the 16S rRNA gene, amplified using the forward primer 338F and the reverse primer 806R. PCR amplification was performed with the MP Taq DNA Polymerase kit (MP Biomedicals, Illkirch, France) under the following thermocycling conditions: an initial denaturation step at 95°C for 5 min, followed by 30 cycles of denaturation at 95°C for 30 s, primer annealing at 52°C for 30 s, and extension at 72°C for 45 s, with a final elongation step at 72°C for 2 min. The resulting amplicons were subjected to paired-end sequencing on an Illumina MiSeq platform (Genotoul sequencing facility, Toulouse, France). Raw sequence quality control and preprocessing were performed using GALAXY FROGS v4.1, resulting in 17356743 high-quality sequences ^78^. Paired-end reads were merged and subsequently clustered into Operational Taxonomic Units (OTUs) using the Swarm v4.1 algorithm. Chimeric sequences and OTUs accounting for less than 0.005% of the total sequence dataset were removed prior to downstream analyses. Taxonomic classification was achieved by querying representative OTU sequences against the SILVA 138 reference database. Read count normalization was performed using DESeq2 v1.38.3, and alpha diversity indices, differential abundance testing, and principal coordinate analysis (PCoA) were performed in the SHAMAN interactive web application ^79^. Functional inference of microbial community metabolic potential was performed using PICRUSt2 (v2.4) applied to FROGS-processed 16S rRNA sequencing data. MetaCyc pathways and enzyme commission (EC) gene families were predicted from taxonomic profiles, with third-tier pathway annotations used to resolve group-specific metabolic differences. Pathway abundances were derived by mapping EC numbers to the MetaCyc database, which was chosen over KEGG for its open-access, community-curated framework, enabling systematic characterization of enzymatic reactions and metabolic networks within each microbial community.

### Untargeted metabolomic analysis

Untargeted metabolomic profiling was carried out at the ENVIROMICS core facility of EBI (UMR CNRS 7267), Université de Poitiers, France. Mouse ceca were emptied, and each sample was stored at –80°C until processing. Approximately 50 mg of homogenized cecal content was weighed into a 2 mL centrifuge tube before the addition of 400 µL of ACN/MeOH/H₂O (2:2:1, v/v/v). Samples were homogenized using a FastPrep-24™ 5G with the recommended “human feces” program (6.0 m/s for 40 s), centrifuged at 14,000 × g for 10 min at 4°C, and 300 µL of the supernatant was transferred to a microtube and stored at –20°C until LC-HRMS analysis. Chromatographic separation was performed on a reversed-phase C18 column (e.g., Waters Acquity BEH C18, 2.1 × 100 mm, 1.7 µm particle size) maintained at 40°C. The mobile phase consisted of ACN/H₂O (5:95, v/v) with 0.1% formic acid (phase A) and ACN/H₂O (95:5, v/v) with 0.1% formic acid (phase B), delivered at 0.3 mL/min using a gradient of 2% B from 0 to 1.5 min, increase to 50% B at 9 min, increase to 95% B at 14 min and held until 17 min, followed by a return to 2% B at 20 min and equilibration until 22 min. The injection volume was set at 10 µL. Mass spectrometry was performed on a Thermo Fisher Scientific Orbitrap Exploris 240 in positive heated electrospray ionization (H-ESI+) mode, with the gas temperature set to 300°C, sheath gas at 50 Arb, auxiliary gas at 10 Arb, sweep gas at 1 Arb, ion transfer tube temperature at 320°C, RF lens at 70%, and capillary voltage at 3500 V. Data were acquired over the m/z range 70–1000 with a cycle time of 0.7 s and a mass tolerance of ± 5 ppm for both MS and MS/MS acquisition. MS scans were acquired at a resolution of 60,000, whereas data-dependent MS² scans were acquired at a resolution of 30,000 with HCD collision energies of 30, 55, and 70%. LC-MS data were acquired using Xcalibur software (Thermo Fisher Scientific, USA) and processed with Compound Discoverer 3.4 (Thermo Fisher Scientific, USA). Peak detection parameters were set to an MS1 mass tolerance of 0.0025 Da and a minimum peak intensity of 30,000, while peak alignment used a retention time tolerance of 0.08 min and an MS1 tolerance of 4 ppm. Compound annotation was performed against mzCloud, MassLists, ChemSpider, and Metabolika databases, followed by manual curation. Data were normalized to the total sum of peak areas. Principal component analysis was performed using MetaboAnalyst 6.0 to assess global variability, and hierarchical clustering heatmaps were generated in MetaboAnalyst 6.0 using auto-scaled features, Euclidean distance, and Ward linkage. Discriminant metabolites were identified by one-way ANOVA across the five experimental groups, followed by Benjamini-Hochberg false discovery rate correction at 5%, and Tukey’s post hoc test was applied to significant features.

### Brain Neurochemical Analysis by 1H Magnetic Resonance Spectroscopy

Brain tissue was collected immediately following sacrifice by rapid decapitation and placed on a pre-cooled dissection plate maintained at –80°C. The cerebral cortex and hippocampus were dissected within 60 seconds of decapitation to prevent post-mortem metabolic alterations, in particular the artifactual accumulation of lactate that occurs progressively following tissue anoxia.

.Neurochemical profiling was performed by 1H High-Resolution Magic Angle Spinning (HR-MAS) NMR spectroscopy on intact frozen tissue biopsies (∼ 40 mg). Samples were loaded into 4 mm zirconia rotors with 10 µL D₂O containing maleate (10 mM) as chemical shift reference. Spectra were acquired on a Bruker Avance III 11.7 T spectrometer (pIBIO, CNRS/Université de Bordeaux UAR 3767) using water suppression sequence (spinning rate: 5,000 Hz; TR = 5 s; 512 transients). Eleven metabolites were quantified: NAA, aspartate, glutamate, glutamine, GABA, alanine, lactate, creatine, myo-inositol, taurine, and choline. Spectra were processed and quantified with a Bruker application.

### Statistical analysis

Data were expressed as mean ± SEM. For body weight gain and physiological test (OGTT) curves, statistical analysis was performed using a two-way repeated-measures analysis of variance (ANOVA) with a post hoc Bonferroni’s test, since time was considered an additional variable (GraphPad Prism, USA). All results were considered statistically significant at p < 0.05.

## Acknowledgments

We thank Laetitia Medan from the University of Bordeaux animal facilities for providing the PILA service for the animals’ care. We acknowledge the Genotoul sequencing platform for providing technical support for 16S rDNA sequencing, and the Genotoul bioinformatics platform and the Sigenae group for providing computing resources. We thank Julien Izotte and Léa MORA CHARROT from Bordeaux University for providing technical support in biochemical analysis. We thank Andrea Boizard Moracchini and Maeva Roy from the CHU de Bordeaux for providing technical support with MSD analysis. We thank Anne Marie Elie and Mathilda Elie from Bordeaux Sciences Agro for providing technical and logistical support. We thank the Service d’Anatomie et Cytologie-Pathologie of CHU de Poitiers.

## Data availability statement

The authors confirm that the data supporting the findings of this study are available within the article and its supplementary materials.

## Ethic statement

Animal experiments were carried out in accordance with the rural and maritime fishing codes R.214-87 to R.214-126 25/04/2025. The Experimental Animal Ethics Committee of Bordeaux University,APAFIS #50103-2024062812454434 v6. The authors have read the ARRIVE guidelines, and the manuscript was prepared in accordance with their recommendations.

## References

1. Ng M, Gakidou E, Lo J, Habtegiorgis Abate Y, Abbafati C, Abbas N, Abbasian M, Abd ElHafeez S, Abdel-Rahman WM, Abd-Elsalam S, et al. Global, regional, and national prevalence of adult overweight and obesity, 1990-2021, with forecasts to 2050: a forecasting study for the Global Burden of Disease Study 2021. The Lancet [Internet] 2025 [cited 2026 Jun 2]; 405:813–38. Available from: https://issp.org/

2. González-Muniesa P, Mártinez-González MA, Hu FB, Després JP, Matsuzawa Y, Loos RJF, Moreno LA, Bray GA, Martinez JA. Obesity. Nature Reviews Disease Primers 2017 3:1 [Internet] 2017 [cited 2026 Jun 2]; 3:17034-. Available from: https://www.nature.com/articles/nrdp201734

3. The Lancet. Childhood obesity: a global health crisis. The Lancet [Internet] 2025 [cited 2026 Jun 2]; 406:1193. Available from: https://pubmed.ncbi.nlm.nih.gov/40975599/

4. Obesity and overweight [Internet]. [cited 2026 Jun 2]; Available from: https://www.who.int/news-room/fact-sheets/detail/obesity-and-overweight

5. Furman D, Campisi J, Verdin E, Carrera-Bastos P, Targ S, Franceschi C, Ferrucci L, Gilroy DW, Fasano A, Miller GW, et al. Chronic inflammation in the etiology of disease across the life span. Nat Med [Internet] 2019 [cited 2026 Jun 2]; 25:1822. Available from: https://pmc.ncbi.nlm.nih.gov/articles/PMC7147972/

6. Soták M, Clark M, Suur BE, Börgeson E. Inflammation and resolution in obesity. Nature Reviews Endocrinology 2024 21:1 [Internet] 2024 [cited 2026 Jun 2]; 21:45–61. Available from: https://www.nature.com/articles/s41574-024-01047-y

7. Tilg H, Zmora N, Adolph TE, Elinav E. The intestinal microbiota fuelling metabolic inflammation. Nat Rev Immunol [Internet] 2020 [cited 2026 Jun 2]; 20:40–54. Available from: https://pubmed.ncbi.nlm.nih.gov/31388093/

8. Zheng D, Liwinski T, Elinav E. Interaction between microbiota and immunity in health and disease. Cell Research 2020 30:6 [Internet] 2020 [cited 2026 Jun 2]; 30:492–506. Available from: https://www.nature.com/articles/s41422-020-0332-7

9. Li Q, de Oliveira Formiga R, Puchois V, Creusot L, Ahmad AH, Amouyal S, Campos-Ribeiro MA, Zhao Y, Harris DMM, Lasserre F, et al. Microbial metabolite indole-3-propionic acid drives mitochondrial respiration in CD4+ T cells to confer protection against intestinal inflammation. Nature Metabolism 2025 7:12 [Internet] 2025 [cited 2026 May 10]; 7:2510–30. Available from: https://www.nature.com/articles/s42255-025-01396-6

10. Chilloux J, Brial F, Everard A, Smyth D, Andrikopoulos P, Zhang L, Plovier H, Myridakis A, Hoyles L, Moreno-Navarrete JM, et al. Inhibition of IRAK4 by microbial trimethylamine blunts metabolic inflammation and ameliorates glycemic control. Nature Metabolism 2025 7:12 [Internet] 2025 [cited 2026 Jun 2]; 7:2531–47. Available from: https://www.nature.com/articles/s42255-025-01413-8

11. Tilg H, Adolph TE, Trauner M. Gut-liver axis: Pathophysiological concepts and clinical implications. Cell Metab [Internet] 2022 [cited 2026 Jun 2]; 34:1700–18. Available from: https://pubmed.ncbi.nlm.nih.gov/36208625/

12. Wu H, Chen J, Guo S, Deng J, Zhou Z, Zhang X, Qi TT, Yu F, Yang Q. Advances in the acting mechanism and treatment of gut microbiota in metabolic dysfunction-associated steatotic liver disease. Gut Microbes [Internet] 2025 [cited 2026 Jun 17]; 17. Available from: https://pubmed.ncbi.nlm.nih.gov/40394806/

13. Loh JS, Mak WQ, Tan LKS, Ng CX, Chan HH, Yeow SH, Foo JB, Ong YS, How CW, Khaw KY. Microbiota–gut–brain axis and its therapeutic applications in neurodegenerative diseases. Signal Transduction and Targeted Therapy 2024 9:1 [Internet] 2024 [cited 2026 Jun 2]; 9:37-. Available from: https://www.nature.com/articles/s41392-024-01743-1

14. Cook TM, Mansuy-Aubert V. Communication between the gut microbiota and peripheral nervous system in health and chronic disease. Gut Microbes [Internet] 2022 [cited 2026 Jun 2]; 14. Available from: https://www.tandfonline.com/doi/pdf/10.1080/19490976.2022.2068365

15. Krüger N, Schneeweiss S, Desai RJ, Sreedhara SK, Kehoe AR, Fuse K, Hahn G, Schunkert H, Wang S V. Cardiovascular outcomes of semaglutide and tirzepatide for patients with type 2 diabetes in clinical practice. Nature Medicine 2026 32:1 [Internet] 2025 [cited 2026 Jun 3]; 32:342–52. Available from: https://www.nature.com/articles/s41591-025-04102-x

16. Lincoff AM, Brown-Frandsen K, Colhoun HM, Deanfield J, Emerson SS, Esbjerg S, Hardt-Lindberg S, Hovingh GK, Kahn SE, Kushner RF, et al. Semaglutide and Cardiovascular Outcomes in Obesity without Diabetes. N Engl J Med [Internet] 2023 [cited 2026 Jun 3]; 389:2221–32. Available from: https://www.nejm.org/doi/pdf/10.1056/NEJMoa2307563

17. Holman RR, Bethel MA, Mentz RJ, Thompson VP, Lokhnygina Y, Buse JB, Chan JC, Choi J, Gustavson SM, Iqbal N, et al. Effects of Once-Weekly Exenatide on Cardiovascular Outcomes in Type 2 Diabetes. N Engl J Med [Internet] 2017 [cited 2026 Jun 3]; 377:1228–39. Available from: https://pubmed.ncbi.nlm.nih.gov/28910237/

18. Gerstein HC, Colhoun HM, Dagenais GR, Diaz R, Lakshmanan M, Pais P, Probstfield J, Riesmeyer JS, Riddle MC, Rydén L, et al. Dulaglutide and cardiovascular outcomes in type 2 diabetes (REWIND): a double-blind, randomised placebo-controlled trial. The Lancet [Internet] 2019 [cited 2026 Jun 3]; 394:121–30. Available from: https://pubmed.ncbi.nlm.nih.gov/31189511/

19. Marso SP, Bain SC, Consoli A, Eliaschewitz FG, Jódar E, Leiter LA, Lingvay I, Rosenstock J, Seufert J, Warren ML, et al. Semaglutide and Cardiovascular Outcomes in Patients with Type 2 Diabetes. N Engl J Med [Internet] 2016 [cited 2026 Jun 3]; 375:1834–44. Available from: https://pubmed.ncbi.nlm.nih.gov/27633186/

20. SP M, GH D, K B-F, P K, JF M, MA N, SE N, S P, NR P, LS R, et al. Liraglutide and Cardiovascular Outcomes in Type 2 Diabetes. N Engl J Med [Internet] 2016 [cited 2026 Jul 29]; 375:101. Available from: https://pubmed.ncbi.nlm.nih.gov/27295427/

21. Daniele G, Campi B, Saba A, Codini S, Ciccarone A, Giusti L, Del Prato S, Esterline RL, Ferrannini E. Plasma N-Acetylaspartate Is Related to Age, Obesity, and Glucose Metabolism: Effects of Antidiabetic Treatment and Bariatric Surgery. Front Endocrinol (Lausanne) [Internet] 2020 [cited 2026 Jun 6]; 11:510676. Available from: www.frontiersin.org

22. Acosta-Estrada BA, Gutiérrez-Uribe JA, Serna-Saldívar SO. Bound phenolics in foods, a review. Food Chem [Internet] 2014 [cited 2024 Dec 4]; 152:46–55. Available from: https://pubmed.ncbi.nlm.nih.gov/24444905/

23. Stiller A, Garrison K, Gurdyumov K, Kenner J, Yasmin F, Yates P, Song BH. From Fighting Critters to Saving Lives: Polyphenols in Plant Defense and Human Health. Int J Mol Sci [Internet] 2021 [cited 2024 Apr 26]; 22. Available from: /pmc/articles/PMC8396434/

24. Mokrani M, Saad N, Nardy L, Sifré E, Despres J, Brochot A, Varon C, Urdaci MC. Biombalance^TM^, an Oligomeric Procyanidins-Enriched Grape Seed Extract, Prevents Inflammation and Microbiota Dysbiosis in a Mice Colitis Model. Antioxidants [Internet] 2025 [cited 2025 Mar 4]; 14:305. Available from: https://www.mdpi.com/2076-3921/14/3/305

25. Mokrani M, Brochot A, Urdaci MC. BiombalanceTM: A Specific Oligomeric Procyanidin-Rich Grape Seed Extract as Multifunctional Ingredient Integrating Antibacterial, Antioxidant, and Anti-Inflammatory Activities with Beneficial Gut–Brain Axis Modulation. Antioxidants [Internet] 2025 [cited 2026 Jan 10]; 14:1484. Available from: https://www.mdpi.com/2076-3921/14/12/1484/htm

26. Crozier A, Jaganath IB, Clifford MN. Dietary phenolics: Chemistry, bioavailability and effects on health. Nat Prod Rep [Internet] 2009 [cited 2025 May 5]; 26:1001–43. Available from: https://pubmed.ncbi.nlm.nih.gov/19636448/

27. Redondo-Castillejo R, Garcimartín A, Hernández-Martín M, López-Oliva ME, Bocanegra A, Macho-González A, Bastida S, Benedí J, Sánchez-Muniz FJ. Proanthocyanidins: Impact on Gut Microbiota and Intestinal Action Mechanisms in the Prevention and Treatment of Metabolic Syndrome. Int J Mol Sci [Internet] 2023 [cited 2026 Jun 17]; 24. Available from: https://pubmed.ncbi.nlm.nih.gov/36982444/

28. Anhê FF, Varin T V., Le Barz M, Desjardins Y, Levy E, Roy D, Marette A. Gut Microbiota Dysbiosis in Obesity-Linked Metabolic Diseases and Prebiotic Potential of Polyphenol-Rich Extracts. Curr Obes Rep 2015; 4:389–400.

29. Mokrani M, Barz M Le, Elie A-M, Renouf É, Mérillon J-M, Limam F, Aouani E, Marette A, Saad N, Urdaci MC. Prevention of Intestinal Inflammation and Gut Dysbiosis by Prebiotic Grape Seed Flour in Mice with DSS-Induced Colitis. Pharmaceuticals 2026, Vol 19, Page 1189 [Internet] 2026 [cited 2026 Jul 31]; 19:1189. Available from: https://www.mdpi.com/1424-8247/19/8/1189/htm

30. Zhao Y, Lu H, Jiang X. Advance in neuroprotective effects of proanthocyanidins (PCs): Structure, absorption, bioactivities, mechanism, and perspectives. Pharmacol Res [Internet] 2026 [cited 2026 Jun 17]; 223. Available from: https://pubmed.ncbi.nlm.nih.gov/41496378/

31. Hu Z, He Z, Wang Y, Chu Z, Zhou Y, Li W, Lu J, Lin Q, Luo F. Targeting the gut-liver axis with dietary polyphenols to ameliorate metabolic dysfunction-associated steatotic liver disease: advances in molecular mechanisms. Crit Rev Food Sci Nutr [Internet] 2026 [cited 2026 Jun 17]; Available from: https://www.tandfonline.com/doi/abs/10.1080/10408398.2025.2556470

32. Zhang Y, Zhou Y, Fang G, Azi F, Gao Z, Wu F, Jiao L, Wang R, Lu B, Liu X. Hepatoprotective potential of anthocyanins from purple highland barley bran against non-alcoholic fatty liver disease through gut-liver axis. Food Res Int [Internet] 2026 [cited 2026 Jun 17]; 230. Available from: https://pubmed.ncbi.nlm.nih.gov/41794448/

33. Arellano-García L, Portillo MP, Hadjihambi A, Martínez JA, Milton-Laskibar I. The Gut–Brain Axis in Obesity: Mechanisms, Development, and Therapeutic Perspectives. Current Nutrition Reports 2026 15:1 [Internet] 2026 [cited 2026 Jun 17]; 15:16-. Available from: https://link.springer.com/article/10.1007/s13668-026-00732-w

34. Yuan C, Wang N, He K, Xu T, Wang H, Ren H, Xue S, Yu Q, Chen L, Zhang G. Dietary Proanthocyanidins Ameliorate Age-Related Cognitive Decline and Neuroinflammation in Mice via the Gut Microbiota-SCFAs-5-HTP Axis. 2026 [cited 2026 Jun 17]; Available from: https://www.researchsquare.com

35. Liu W, Zhao S, Wang J, Shi J, Sun Y, Wang W, Ning G, Hong J, Liu R. Grape seed proanthocyanidin extract ameliorates inflammation and adiposity by modulating gut microbiota in high-fat diet mice. Mol Nutr Food Res [Internet] 2017 [cited 2024 Dec 4]; 61. Available from: https://pubmed.ncbi.nlm.nih.gov/28500724/

36. Liu M, Yun P, Hu Y, Yang J, Khadka RB, Peng X. Effects of Grape Seed Proanthocyanidin Extract on Obesity. Obes Facts [Internet] 2020 [cited 2026 Jun 15]; 13:279–91. Available from: https://pubmed.ncbi.nlm.nih.gov/32114568/

37. Baron G, Altomare A, Della Vedova L, Gado F, Quagliano O, Casati S, Tosi N, Bresciani L, Del Rio D, Roda G, et al. Unraveling the parahormetic mechanism underlying the health-protecting effects of grapeseed procyanidins. Redox Biol [Internet] 2024 [cited 2026 Jun 15]; 69:102981. Available from: https://www.sciencedirect.com/science/article/pii/S2213231723003828

38. Cawthorn WP, Sethi JK. TNF-α and adipocyte biology. FEBS Lett [Internet] 2007 [cited 2026 Jun 15]; 582:117. Available from: https://pmc.ncbi.nlm.nih.gov/articles/PMC4304634/

39. Darkoh C, Chappell C, Gonzales C, Okhuysen P. A rapid and specific method for the detection of indole in complex biological samples. Appl Environ Microbiol [Internet] 2015 [cited 2026 May 21]; 81:8093–7. Available from: https://pubmed.ncbi.nlm.nih.gov/26386049/

40. Broskey NT, Zou K, Dohm GL, Houmard JA. Plasma Lactate as a Marker for Metabolic Health. Exerc Sport Sci Rev [Internet] 2020 [cited 2026 Jun 16]; 48:119. Available from: https://pmc.ncbi.nlm.nih.gov/articles/PMC7311283/

41. Mokrani M, Charradi K, Limam F, Aouani E, Urdaci MC. Grape seed and skin extract, a potential prebiotic with anti – obesity effect through gut microbiota modulation. Gut Pathog [Internet] 2022;:1–14. Available from: 10.1186/s13099-022-00505-0

42. Ravnskjaer K, Frigerio F, Boergesen M, Nielsen T, Maechler P, Mandrup S. PPARδ is a fatty acid sensor that enhances mitochondrial oxidation in insulin-secreting cells and protects against fatty acid-induced dysfunction. J Lipid Res [Internet] 2010 [cited 2026 Jun 16]; 51:1370. Available from: https://pmc.ncbi.nlm.nih.gov/articles/PMC3035500/

43. Herman MA, Samuel VT. The Sweet Path to Metabolic Demise: Fructose and Lipid Synthesis. Trends in Endocrinology and Metabolism [Internet] 2016 [cited 2026 Jun 16]; 27:719–30. Available from: https://pubmed.ncbi.nlm.nih.gov/27387598/

44. Shao W, Espenshade PJ. Expanding roles for SREBP in metabolism. Cell Metab [Internet] 2012 [cited 2026 Jun 16]; 16:414. Available from: https://pmc.ncbi.nlm.nih.gov/articles/PMC3466394/

45. Osborn LJ, Schultz K, Massey W, DeLucia B, Choucair I, Varadharajan V, Banerjee R, Fung K, Horak AJ, Orabi D, et al. A gut microbial metabolite of dietary polyphenols reverses obesity-driven hepatic steatosis. Proc Natl Acad Sci U S A [Internet] 2022 [cited 2026 Jun 16]; 119:e2202934119. Available from: 10.1073/pnas.2202934119?download=true

46. Wan Y, Yuan J, Li J, Li H, Zhang J, Tang J, Ni Y, Huang T, Wang F, Zhao F, et al. Unconjugated and secondary bile acid profiles in response to higher-fat, lower-carbohydrate diet and associated with related gut microbiota: A 6-month randomized controlled-feeding trial. Clinical Nutrition [Internet] 2020 [cited 2026 Jul 31]; 39:395–404. Available from: https://pubmed.ncbi.nlm.nih.gov/30876827/

47. Park S, Zhang T, Yue Y, Wu X. Effects of Bile Acid Modulation by Dietary Fat, Cholecystectomy, and Bile Acid Sequestrant on Energy, Glucose, and Lipid Metabolism and Gut Microbiota in Mice. Int J Mol Sci [Internet] 2022 [cited 2026 Jul 31]; 23:5935. Available from: https://pmc.ncbi.nlm.nih.gov/articles/PMC9180239/

48. Iliev ID, Ananthakrishnan AN, Guo CJ. Microbiota in inflammatory bowel disease: mechanisms of disease and therapeutic opportunities. Nature Reviews Microbiology 2025 23:8 [Internet] 2025 [cited 2026 Jun 16]; 23:509–24. Available from: https://www.nature.com/articles/s41579-025-01163-0

49. Holst JJ. The physiology of glucagon-like peptide 1. Physiol Rev [Internet] 2007 [cited 2026 Jun 15]; 87:1409–39. Available from: https://journals.physiology.org/doi/pdf/10.1152/physrev.00034.2006?download=true

50. Ranganath LR, Beety JM, Morgan LM, Wright JW, Howland R, Marks V. Attenuated GLP-1 secretion in obesity: cause or consequence? Gut [Internet] 1996 [cited 2026 Jun 15]; 38:916. Available from: https://pmc.ncbi.nlm.nih.gov/articles/PMC1383202/

51. Yoon HS, Cho CH, Yun MS, Jang SJ, You HJ, Kim J hyeong, Han D, Cha KH, Moon SH, Lee K, et al. Akkermansia muciniphila secretes a glucagon-like peptide-1-inducing protein that improves glucose homeostasis and ameliorates metabolic disease in mice. Nat Microbiol [Internet] 2021 [cited 2026 Jun 15]; 6:563–73. Available from: 10.1038/s41564-021-00880-5

52. Suenaert P, Segers A, Rymenans L, Devroye H, Moll JM, Cani PD, de Vos WM. Effect of pasteurized Akkermansia muciniphila MucT on insulin sensitivity, body composition, and GLP-1 production in subjects with metabolic syndrome: impact of low baseline gut Akkermansia levels. Gut Microbes [Internet] 2026 [cited 2026 Jul 31]; 18. Available from: https://www.tandfonline.com/doi/pdf/10.1080/19490976.2026.2690689

53. Irfan Z, Halder J, Giri S, Molla EA, Khanam S. Therapeutic potential of prebiotics in modulating postprandial GLP-1, GLP-2, and glucose homeostasis in type 2 diabetes mellitus: Targeting gut dysbiosis and insulin resistance. Diabetes Res Clin Pract [Internet] 2026 [cited 2026 Jun 15]; 232:113102. Available from: https://www.sciencedirect.com/science/article/pii/S0168822726000215

54. Kamath S, Chan NSL, Joyce P. GLP-1 agonists and the gut microbiome: A bidirectional relationship. Br J Clin Pharmacol [Internet] 2026 [cited 2026 Jun 15]; 92. Available from: https://pubmed.ncbi.nlm.nih.gov/41703894/

55. Cryan JF, O’riordan KJ, Cowan CSM, Sandhu K V., Bastiaanssen TFS, Boehme M, Codagnone MG, Cussotto S, Fulling C, Golubeva A V., et al. The Microbiota-Gut-Brain Axis. Physiol Rev [Internet] 2019 [cited 2026 Jun 15]; 99:1877–2013. Available from: https://pubmed.ncbi.nlm.nih.gov/31460832/

56. Mendoza-Herrera K, Florio AA, Moore M, Marrero A, Tamez M, Bhupathiraju SN, Mattei J. The Leptin System and Diet: A Mini Review of the Current Evidence. Front Endocrinol (Lausanne) [Internet] 2021 [cited 2026 Jun 15]; 12:749050. Available from: https://pmc.ncbi.nlm.nih.gov/articles/PMC8651558/

57. Strandwitz P. Neurotransmitter modulation by the gut microbiota. Brain Res [Internet] 2018 [cited 2026 Jun 15]; 1693:128–33. Available from: https://pubmed.ncbi.nlm.nih.gov/29903615/

58. Yang M, Bose S, Lim S, Seo J, Shin J, Lee D, Chung WH, Song EJ, Nam Y Do, Kim H. Beneficial effects of newly isolated Akkermansia muciniphila strains from the human gut on obesity and metabolic dysregulation. Microorganisms 2020; 8:1413.

59. Paone P, Petitfils C, Puel A, Latousakis D, de Vos WM, Delzenne NM, Juge N, Van Hul M, Cani PD. Akkermansia muciniphila modulates intestinal mucus composition to counteract high-fat diet-induced obesity in mice. Gut Microbes [Internet] 2026 [cited 2026 Jun 16]; 18:2612580. Available from: https://www.tandfonline.com/doi/pdf/10.1080/19490976.2025.2612580

60. Depommier C, Everard A, Druart C, Plovier H, Van Hul M, Vieira-Silva S, Falony G, Raes J, Maiter D, Delzenne NM, et al. Supplementation with Akkermansia muciniphila in overweight and obese human volunteers: a proof-of-concept exploratory study. Nat Med [Internet] 2019 [cited 2026 Jun 16]; 25:1096–103. Available from: https://pubmed.ncbi.nlm.nih.gov/31263284/

61. Mount S, Canfora EE, Jocken JW, Umanets A, Hul G, Coenjaerds M, Aldaz Laquidain P, Adriaens ME, Holst JJ, Jardon KM, et al. Pasteurized Akkermansia muciniphila MucT for weight loss maintenance in people with overweight and obesity: a controlled randomized trial. Nature Medicine 2026 [Internet] 2026 [cited 2026 Jun 16];:1–10. Available from: https://www.nature.com/articles/s41591-026-04394-7

62. Anhê FF. A polyphenol-rich cranberry extract protects from diet-induced obesity, insulin resistance and intestinal inflammation in association with increased Akkermansia spp. population in the gut microbiota of mice. Gut 64:872.

63. Zhang L, Carmody RN, Kalariya HM, Duran RM, Moskal K, Poulev A, Kuhn P, Tveter KM, Turnbaugh PJ, Raskin I, et al. Grape proanthocyanidin-induced intestinal bloom of Akkermansia muciniphila is dependent on its baseline abundance and precedes activation of host genes related to metabolic health. J Nutr Biochem [Internet] 2018 [cited 2026 Jun 16]; 56:142–51. Available from: https://www.sciencedirect.com/science/article/pii/S0955286317310537?via%3Dihub

64. Cani PD, Depommier C, Derrien M, Everard A, de Vos WM. Akkermansia muciniphila: paradigm for next-generation beneficial microorganisms. Nature Reviews Gastroenterology & Hepatology 2022 [Internet] 2022 [cited 2022 Jun 18];:1–13. Available from: https://www.nature.com/articles/s41575-022-00631-9

65. Rodríguez-Daza MC, Boeren S, Tytgat HLP, Desjardins Y, de Vos WM. Akkermansia muciniphila MucT harnesses dietary polyphenols as xenosiderophores for enhanced iron uptake. Nature Communications 2025 16:1 [Internet] 2025 [cited 2026 Feb 11]; 16:9428-. Available from: https://www.nature.com/articles/s41467-025-64477-w

66. Pabón ML, Lönnerdal B. Effects of type of fat in the diet on iron bioavailability assessed in suckling and weanling rats. Journal of Trace Elements in Medicine and Biology [Internet] 2001 [cited 2026 Jun 16]; 15:18–23. Available from: https://www.sciencedirect.com/science/article/pii/S0946672X01800213

67. Guevara Agudelo FA, Leblanc N, Bourdeau-Julien I, St-Arnaud G, Dahhani F, Flamand N, Veilleux A, Marzo V Di, Raymond F. Dietary iron interacts with diet composition to modulate the endocannabinoidome and the gut microbiome in mice.

68. Chen X, Li Q, Zhang W, Xu Y, Nie X, Wang X, Chen C, Xie J, Nie S. Akkermansia muciniphila-derived extracellular vesicles alleviate colitis-related cognitive impairment via tryptophan metabolic reprogramming of the gut brain axis. Gut Microbes [Internet] 2026 [cited 2026 Jun 16]; 18. Available from: https://www.tandfonline.com/doi/pdf/10.1080/19490976.2025.2611546

69. Kang EJ, Cha MG, Kwon GH, Han SH, Yoon SJ, Lee SK, Ahn ME, Won SM, Ahn EH, Suk KT. Akkermansia muciniphila improve cognitive dysfunction by regulating BDNF and serotonin pathway in gut-liver-brain axis. Microbiome 2024 12:1 [Internet] 2024 [cited 2026 Jun 16]; 12:181-. Available from: https://link.springer.com/article/10.1186/s40168-024-01924-8

70. Moffett JR, Ross B, Arun P, Madhavarao CN, Namboodiri AMA. N-Acetylaspartate in the CNS: From neurodiagnostics to neurobiology. Prog Neurobiol [Internet] 2007 [cited 2026 Jun 16]; 81:89–131. Available from: https://pubmed.ncbi.nlm.nih.gov/17275978/

71. Vuković M, Nosek I, Slotboom J, Medić Stojanoska M, Kozić D. Neurometabolic Profile in Obese Patients: A Cerebral Multi-Voxel Magnetic Resonance Spectroscopy Study. Medicina 2024, Vol 60, Page 1880 [Internet] 2024 [cited 2026 Jun 16]; 60:1880. Available from: https://www.mdpi.com/1648-9144/60/11/1880/htm

72. Li H, Fang Y, Wang D, Shi B, Thompson GJ. Impaired brain glucose metabolism in glucagon-like peptide-1 receptor knockout mice. Nutrition & Diabetes 2024 14:1 [Internet] 2024 [cited 2026 Jul 30]; 14:86-. Available from: https://www.nature.com/articles/s41387-024-00343-w

73. Roy M, Beauvieux MC, Naulin J, El Hamrani D, Gallis JL, Cunnane SC, Bouzier-Sore AK. Rapid adaptation of rat brain and liver metabolism to a ketogenic diet: an integrated study using (1)H– and (13)C-NMR spectroscopy. J Cereb Blood Flow Metab [Internet] 2015 [cited 2026 Jul 26]; 35:1154–62. Available from: https://pubmed.ncbi.nlm.nih.gov/25785828/

74. Sears SMS, Hewett SJ. Influence of glutamate and GABA transport on brain excitatory/inhibitory balance. Exp Biol Med [Internet] 2021 [cited 2026 Jul 26]; 246:1069–83. Available from: https://pubmed.ncbi.nlm.nih.gov/33554649/

75. Anhê FF, Roy D, Pilon G, Dudonné S, Matamoros S, Varin T V., Garofalo C, Moine Q, Desjardins Y, Levy E, et al. A polyphenol-rich cranberry extract protects from diet–induced obesity, insulin resistance and intestinal inflammation in association with increased Akkermansia spp. population in the gut microbiota of mice. Gut [Internet] 2015 [cited 2026 Jun 3]; 64:872–83. Available from: https://pubmed.ncbi.nlm.nih.gov/25080446/

76. Kleiner DE, Brunt EM, Van Natta M, Behling C, Contos MJ, Cummings OW, Ferrell LD, Liu YC, Torbenson MS, Unalp-Arida A, et al. Design and validation of a histological scoring system for nonalcoholic fatty liver disease. Hepatology [Internet] 2005 [cited 2026 Jun 20]; 41:1313–21. Available from: https://pubmed.ncbi.nlm.nih.gov/15915461/

77. Mokrani M, Jacquot C. ENCAPSULATION OF A PEDIOCIN PA-1 PRODUCER PEDIOCOCCUS ACIDILACTICI AND ITS IMPACT ON ENHANCED SURVIVAL AND GUT MICROBIOTA MODULATION. 2024 [cited 2024 Dec 23]; Available from: https://www.researchgate.net/publication/381796554

78. Escudié F, Auer L, Bernard M, Mariadassou M, Cauquil L, Vidal K, Maman S, Hernandez-Raquet G, Combes S, Pascal G. FROGS: Find, Rapidly, OTUs with Galaxy Solution. Bioinformatics 2018; 34:1287–94.

79. Volant S, Lechat P, Woringer P, Motreff L, Campagne P, Malabat C, Kennedy S, Ghozlane A. SHAMAN: A user-friendly website for metataxonomic analysis from raw reads to statistical analysis. BMC Bioinformatics 2020; 21:1–15.

